# A neural and behavioral tradeoff underlies exploratory decisions in normative anxiety

**DOI:** 10.1101/2020.09.16.300525

**Authors:** Kristoffer C. Aberg, Ido Toren, Rony Paz

**Author notes:** **Correspondence** should be addressed to: K.A. or R.P.

## Abstract

Exploration reduces uncertainty about the environment and improves the quality of future decisions, but at the cost of provisional uncertain and suboptimal outcomes. Although anxiety promotes intolerance to uncertainty, it remains unclear whether and by which mechanisms anxiety modulates exploratory decision-making. We use a dynamic three-armed-bandit task and find that higher trait-anxiety increases exploration, which in turn harms overall performance. We identify two distinct behavioral sources: first, decisions made by anxious individuals were guided towards reduction of uncertainty; and second, decisions were less guided by immediate value gains. Imaging (fMRI) revealed that anxiety correlated negatively with the representation of expected-value in the dorsal-anterior-cingulate-cortex, and in contrast, positively with the representation of uncertainty in the anterior-insula. Moreover, the findings were similar in both loss and gain domains, and demonstrated that affective trait modulates exploration and results in an inverse-U-shaped relationship between anxiety and overall performance. We conclude that a shift in balance towards representation of uncertainty in the insula prevails over reduced value representation in the dACC, and entails maladaptive decision-making in individuals with higher normal-range anxiety.

## Introduction

Exploratory decision-making concerns the balance between exploiting options with known outcomes, such as dining in your favorite restaurant; and exploring options with uncertain but potentially better outcomes, such as sampling a novel restaurant. Exploration provides information about the environment and therefore provides a prospective benefit to decision-making by reducing the uncertainty associated with available options. For example, the benefits and costs of eating at a new restaurant are largely unknown until it has been explored, at which point the newly gathered information can be used to guide subsequent decisions (whether to return or not). Exploratory decisions are prevalent in everyday life and involve decisions ranging from small to large and even potentially life-changing ones, as in selecting a partner or employment (Cohen, McClure, and Yu 2007; Addicott et al. 2017; Mehlhorn et al. 2015). Because exaggerated exploration impairs the ability to exploit the good familiar options, whereas overly restrained exploration prevents the discovery of better options, the ability to maintain an appropriate balance has a significant influence on an individual’s well-being and quality of life. In extreme cases, it might even contribute to psychopathologies (Addicott et al. 2017; Scholl and Klein-Flugge 2018).

Imaging studies in humans have shown that exploratory decisions engage the anterior insulae (aINS) as well as prefrontal regions involved in executive control, such as the frontopolar cortex (FPC) and the dorsal anterior cingulate cortex (dACC) (Daw et al. 2006; Badre et al. 2012; Cavanagh et al. 2012; Kayser et al. 2015; Laureiro-Martinez et al. 2015; Blanchard and Gershman 2018; Chakroun et al. 2020; Tomov et al. 2020), with recent research suggesting further differentiation between neuronal correlates related to ‘directed’ and ‘random’ exploration strategies (Tomov et al. 2020). Importantly, it was shown that anxious individuals find uncertainty more aversive and are also more intolerant thereof (Hartley and Phelps 2012; Buhr and Dugas 2009; Grupe and Nitschke 2013; Maner and Schmidt 2006; Maner et al. 2007; Charpentier et al. 2017), and recent studies have shown that aspects of uncertainty correlate with a hyper-responsive aINS in anxiety (Grupe and Nitschke 2013; Paulus and Stein 2006). In addition, anxiety also pertain to dysfunctional executive control processes in the prefrontal-cortex (PFC) which are hypothesized to induce attentional biases towards salient events (Etkin and Wager 2007; Bishop 2007).

These studies suggest that anxiety may have a particularly relevant influence on exploratory decision-making. On one hand, anxiety could decrease exploration because it requires selecting options with uncertain outcomes; yet on the other hand, anxiety could increase exploration by reducing uncertainty in the environment. Previous studies have examined the relationship between uncertainty and anxiety (Grupe and Nitschke 2013) and separately examined the relationship between uncertainty and exploration (Badre et al. 2012; Cavanagh et al. 2012; Kayser et al. 2015; Tomov et al. 2020). However, little is known about the direct relationship between anxiety and exploratory decision-making, the cognitive factors that contribute to it, and the underlying brain mechanisms. Moreover, although outcome valence (appetitive – aversive) was shown to interact with inter-individual factors and anxiety levels (Browning et al. 2015; Laufer, Israeli, and Paz 2016; Xu et al. 2013; Aberg, Doell, and Schwartz 2015, 2016b; Frank, Seeberger, and O’Reilly R 2004; Aberg, Muller, and Schwartz 2017), and activity in the dACC and aINS (Skvortsova, Palminteri, and Pessiglione 2014; Pessiglione and Delgado 2015), little is known about the interaction between these factors and exploratory decisions.

Taken together, we hypothesized that anxiety-levels would induce a bias in exploratory decisions, and that it can be mediated by shifts in saliency to uncertainty and to expected-outcomes, induced by a hyper-responsive aINS and/or by low recruitment of the FPC/dACC.

## Results

Participants played a three-armed bandit task where in each trial they chose one of three slot machines (Fig.1A,B). After selection, the outcome currently associated with the selected machine was displayed. Exploration was promoted by inducing a dynamic environment in which the outcome associated with each machine varied across trials: each arm was associated with a sine-like wave of similar magnitude and mean, but with different period lengths (100, 67, or 33 trials), and phase-shifts (periods of 0.10, 0.45, and 0.90; Fig.1C). To examine differences in exploratory strategies between aversive and appetitive contexts, participants either accumulated points in a Gain condition where all outcomes are positive (Fig.1A), or avoided losing points in a Loss condition where all outcomes are negative (Fig.1B).

**Figure 1.**
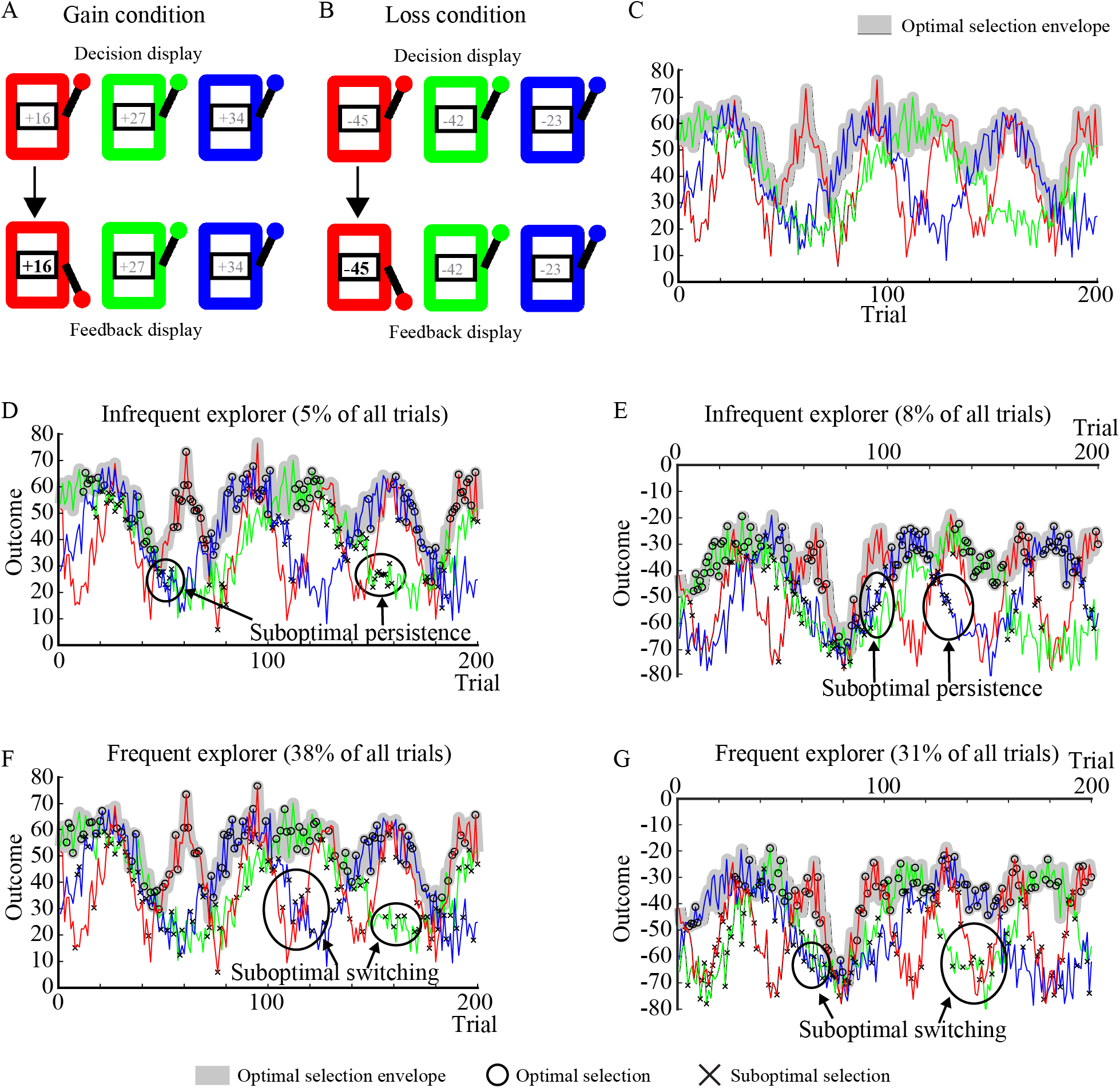
Paradigm and individual behavior. **A.** Illustration of Decision and Feedback displays in the Gain condition, where all outcomes are positive. During the decision display, the outcomes of each machine is hidden (grey characters), and participants decide to play one of the three slot machines by pressing the corresponding button. After selecting a slot machine its outcome is revealed (+16 points), while the outcomes of the non-selected machines remain hidden. **B.** In the Loss condition, all outcomes are negative and participants aim to avoid large losses. **C.** Exploration is promoted by varying the outcome associated with each machine across trials. The optimal potential outcome is displayed in grey. **D,E.** Examples of infrequent explorers in the Gain (**D**) and in the Loss (**E**) condition. **F,G.** Examples of frequent explorers in the Gain (**F**) and in the Loss (**G**) condition.

We first performed a separate behavioral study to test the hypothesis, and found a confirmative positive correlation between trait anxiety and exploration (Supplementary Note 1). Following this, we conducted the independent main behavioral-fMRI study, and examined the interaction across the range of anxiety (Trait-anxiety scores: mean=36.321, SD=9.982; see (Spielberger et al. 1983), Table 1). Normative anxiety is a major factor in modern lives affecting daily function, and enabled us to examine the gradual modulation of anxiety on exploration without the traditional focus and confounds imposed by clinical populations.

### Exploration increases with anxiety

We examined the relationship between Trait anxiety and exploratory decision-making, and whether it interacts with outcome valence (Loss/Gain). As in previous studies (Tzovara et al. 2012; Daw et al. 2006), an exploratory decision was defined as the selection of a machine with a suboptimal expected value, if the selected machine was also different from the machine selected in the previous trial. Trial-by-trial estimates of expected values were estimated by a behavioral model which provided the most parsimonious fit to overall-behavior, as determined by confronting different types of behavioral models commonly used to model behavior and exploration in multi-armed bandit tasks (see below, Methods, and Supplementary Note 5). Infrequent explorers persevered in selecting a suboptimal machine (Fig.1D,E; *suboptimal persistence*), whereas frequent explorers more often rejected the optimal machine and chose a different one (Fig.1F,G; *suboptimal switching*).

We found that exploration, quantified as the proportion of exploratory decisions across all trials, was modulated by anxiety (Fig.2A; repeated measures two-way ANOVA, main effect of anxiety as linear covariate, F(1,26)=24.98, p<0.001, 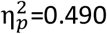, ANOVA; Loss: r=0.707, p<0.001; Gain: r=0.573, p<0.001, two-tailed Pearson correlations; no valence*anxiety interaction effect, F(1,26)=1.85, p=0.19, 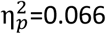, ANOVA). Therefore, more anxious individuals explored more.

**Figure 2.**
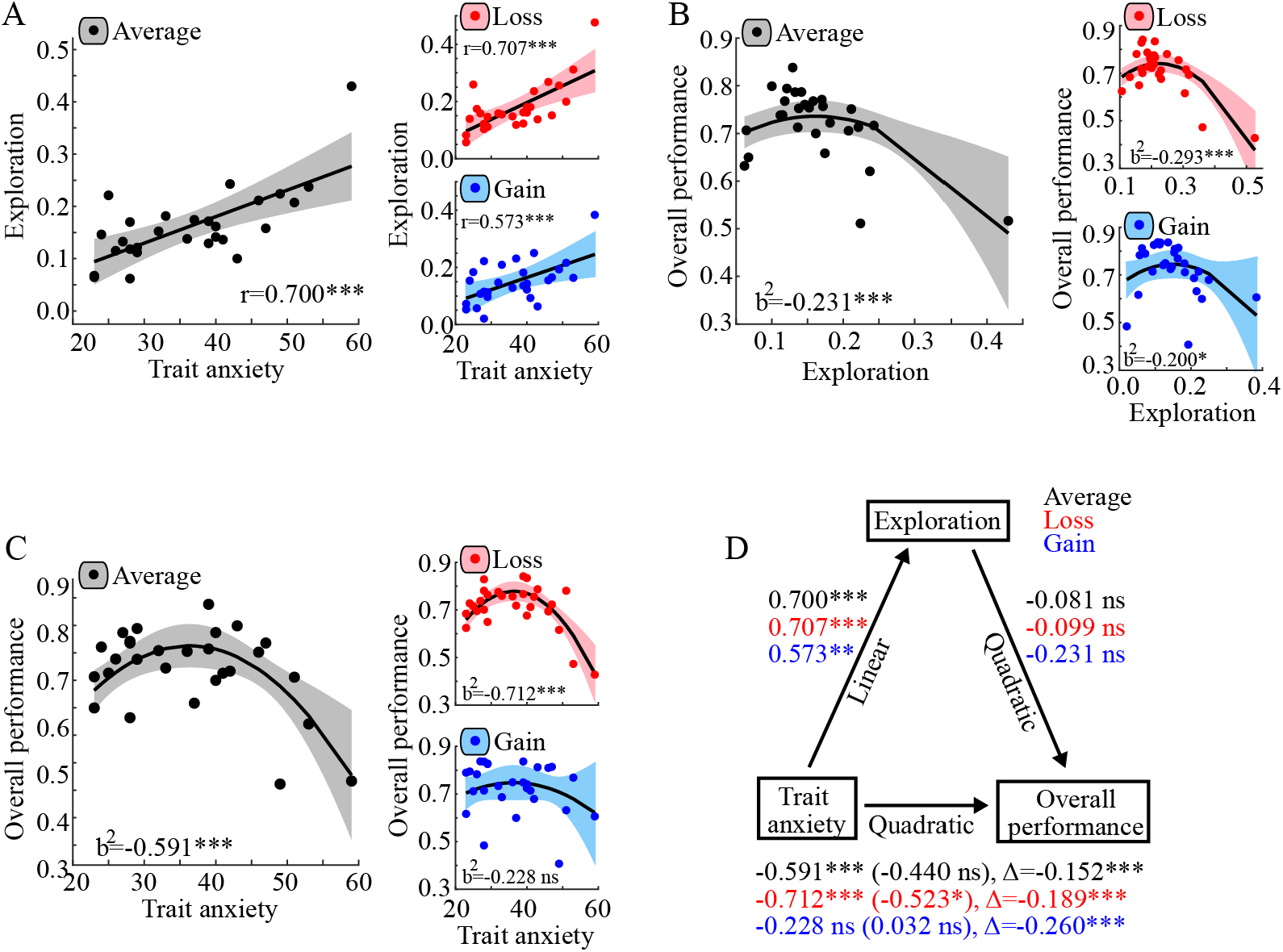
Behavioral results. **A.** Exploration, the proportion of exploratory decisions, increased linearly with increased levels of trait anxiety across conditions (left panel), and separately in the Loss (top right panel) and in the Gain (bottom right panel) condition. **B.** Overall performance, defined as the correlation between maximal and actual outcomes across all trials, varied non-linearly with exploration across conditions (left panel), and separately in the Loss (top right panel) and in the Gain condition (bottom right panel). **C.** Overall performance varied significantly and non-linearly with trait anxiety across conditions (left panel), and in the Loss (top right panel), but not significantly so in the Gain condition (bottom right panel). **D.** Standardized regression coefficients for the relationships between trait anxiety and overall performance as mediated by exploration. The standardized regression coefficient between trait anxiety and overall performance, when controlling for exploration, is in parenthesis. The relationship between trait anxiety and overall performance was mediated by exploration across conditions (black text), as well as in both the Loss (red text) and the Gain condition (blue text), as indicated by significantly reduced quadratic regression coefficients when controlling for exploration (both p-values <0.001). ***p<0.001; *p<0.05; ns p>0.05.

### Overall performance is inverse-U-shape modulated by exploration

In a framework of varying outcomes, it is expected that low exploration would prevent the discovery of changes and better alternatives (Infrequent explorers; Fig.1D,E), and on the other hand, high exploration would hinder the exploitation of the current optimal alternative (Frequent explorers; Fig.1F,G). Taken together, this should lead to an inverse-U-shape form of overall performance.

Indeed, both infrequent and frequent explorers made more suboptimal decisions, and as a result overall performance was non-linearly modulated by the proportion of exploratory decisions (Fig.2C). This was the case when data was collapsed across conditions (Fig.2B, left panel; nonlinear regression with intercept and standardized quadratic term b^2^_Explore_=−0.231, p<0.001), and separately in both the Loss condition (Fig.2B, top right panel; nonlinear regression with intercept and standardized quadratic term b^2^_Explore_=−0.293, p<0.001) and in the Gain condition (Fig.2B, bottom right panel; standardized b^2^_Explore_=−0.200, p<0.05). There was no significant difference between conditions (Actual difference: b^2^_Explore_ Loss-Gain=−0.093; Null distribution difference: b^2^_Explore_ Loss-Gain=−0.040, p=0.198, Monte-Carlo randomization test, Supplementary Note 2). Therefore, our findings demonstrate that the individual tendency to explore modulates overall performance in an inverted U-shape fashion, as previously suggested (Addicott et al. 2017; Aston-Jones and Cohen 2005).

### Exploration mediates the modulation of overall performance by anxiety

If exploration increases with anxiety and overall performance is non-linearly modulated by exploration, then one can hypothesize a relationship between anxiety and performance. We tested this directly and found that overall performance was also significantly and non-linearly modulated by anxiety across conditions (Fig.2C, left panel, nonlinear regression with intercept and standardized quadratic term b^2^_Anxiety_=−0.591, p<0.001), but only significantly so in the Loss condition (Fig.2C, top right panel, nonlinear regression with intercept and standardized quadratic term b^2^_Anxiety_=−0.712, p<0.001), and not in the Gain condition (Fig.2 C, bottom right panel, standardized b^2^_Anxiety_=−0.228, p=0.198), with a significant difference between conditions (Actual difference: b^2^_Anxiety_ Loss-Gain=−0.483; Null distribution difference: b^2^_Anxiety_ Loss-Gain=−0.258, p=0.032, Monte-Carlo permutation test, Supplementary Note 3).

To establish further that it is the exploration that mediated the relationship between anxiety and overall performance, we tested whether the relationship between anxiety and performance is reduced when controlling for the contribution of exploration (Hayes and Rockwood 2017). The relationship changed significantly when controlling for Exploration across conditions (Δb^2^_Anxiety_=−0.190, p<0.001, Monte-Carlo randomization test), and separately in both the Loss (Δb^2^_Anxiety_=−0.152, p<0.001, Monte-Carlo randomization test) and in the Gain condition (Δb^2^_Anxiety_=−0.260, p<0.001, Monte-Carlo randomization test; see Fig.2D and Supplementary Note 4 for details). Therefore, exploration contributes to the non-linear relationship between anxiety and overall performance.

### Behavioral factors contributing to exploration

We examined the relative contribution of different behavioral factors by adapting a reinforcement-learning approach. These models track the expected-value of each machine and update it by the mismatch between the expected-value and the actual outcome (i.e. the prediction-error scaled by the learning rate. Methods, Equations 1,2,4). In addition, we included the following factors: First, because recent evidence suggests that exploration is aimed at reducing the outcome-variability associated with different options (‘directed’ exploration in static environments: (Badre et al. 2012; Frank et al. 2009; Tomov et al. 2020; Gershman 2018), we included an outcome-variability factor which was updated by the unsigned prediction-error scaled by a learning-rate (Equations 5–6). Second, in a dynamic environment the uncertainty associated with a machine’s outcome increases with the time since it was last selected, and we therefore included an outcome-uncertainty factor as an additional way of modeling ‘directed’ exploration, defined as the number of trials since a machine was last selected (Equation 7). Third, an exploratory strategy that allows detecting changes in the environment is randomly switching machines between trials, and to capture such tendencies we included a random binary switch factor (Equation 8). Notice this is the inverse of ‘stickiness’ or ‘perseveration’, which models the tendency to repeat a previously chosen action (Chakroun et al. 2020; Gershman, Pesaran, and Daw 2009). Finally, ‘random’ exploration in static environments was found to depend on the total uncertainty of the environment (Gershman 2018; Gershman and Tzovaras 2018), and we therefore included a term which scales the expected-value of each machine by the total outcome-variability of the environment (Equation 9).

In addition to the models described above, we also tested two common models: one with an ε-greedy decision rule in which suboptimal alternatives were selected with probability ε (Equation 11), as well as Kalman-filter model with dynamic learning rates that was previously used to disentangle ‘directed’ and ‘random’ exploration in static environments (Equations 12–15).

After the models were fitted individually to each participant’s behavior, the probability of selecting machine *i* in trial *t* depends on weights β_*Q*_, β_*U*_, β_*T*_, β_*S*_, and β_*tU*_ reflecting the contribution of expected-value *Q*_*i*_(*t*), outcome-variability *U*_*i*_(*t*), outcome-uncertainty *T*_*i*_(*t*), random-switching *S*_*i*_(*t*), and ‘random exploration’ *Q*_*i*_(*t*)/*tU*_*i*_(*t*), respectively. The goodness-of-fit of these models were compared using a Bayesian Model Selection (BMS) procedure (Stephan et al. 2009), with Akaike’s Information Criterion corrected for small sample sizes as model evidence (AICc; (Burnham and Anderson 2004)), and further validated using the Bayesian Information Criterion (BIC) approximation of the log marginal likelihood (Bishop 2006; Gershman 2018), which provides a larger penalty term for more complex models.

We found that the most parsimonious model was the same when collapsed across valence conditions (Fig.3A, protected exceedance probabilities > 0.997; Supplementary Note 5) as well as for each loss/gain condition separately (Fig.3B). This model included decision weights for expected-value (β_*Q*_), outcome-uncertainty (β_*T*_), and random-switching (β_*S*_) and it is therefore termed the QTS model hereafter. Importantly, the QTS model was found to be the most parsimonious one also in the independent behavioral pilot study (Supplementary Note 1). Next, to ensure that the fitted parameters were meaningful, we applied a parameter recovery procedure which tests whether model-parameters used to generate behavior can be successfully recovered when re-fitting the same model to the generated behavior (Wilson and Collins 2019). Indeed, all correlations between generating and recovered parameters were highly significant (Fig.3C; all two-sided Pearson’s r>0.827, p-values<0.001).

**Figure 3.**
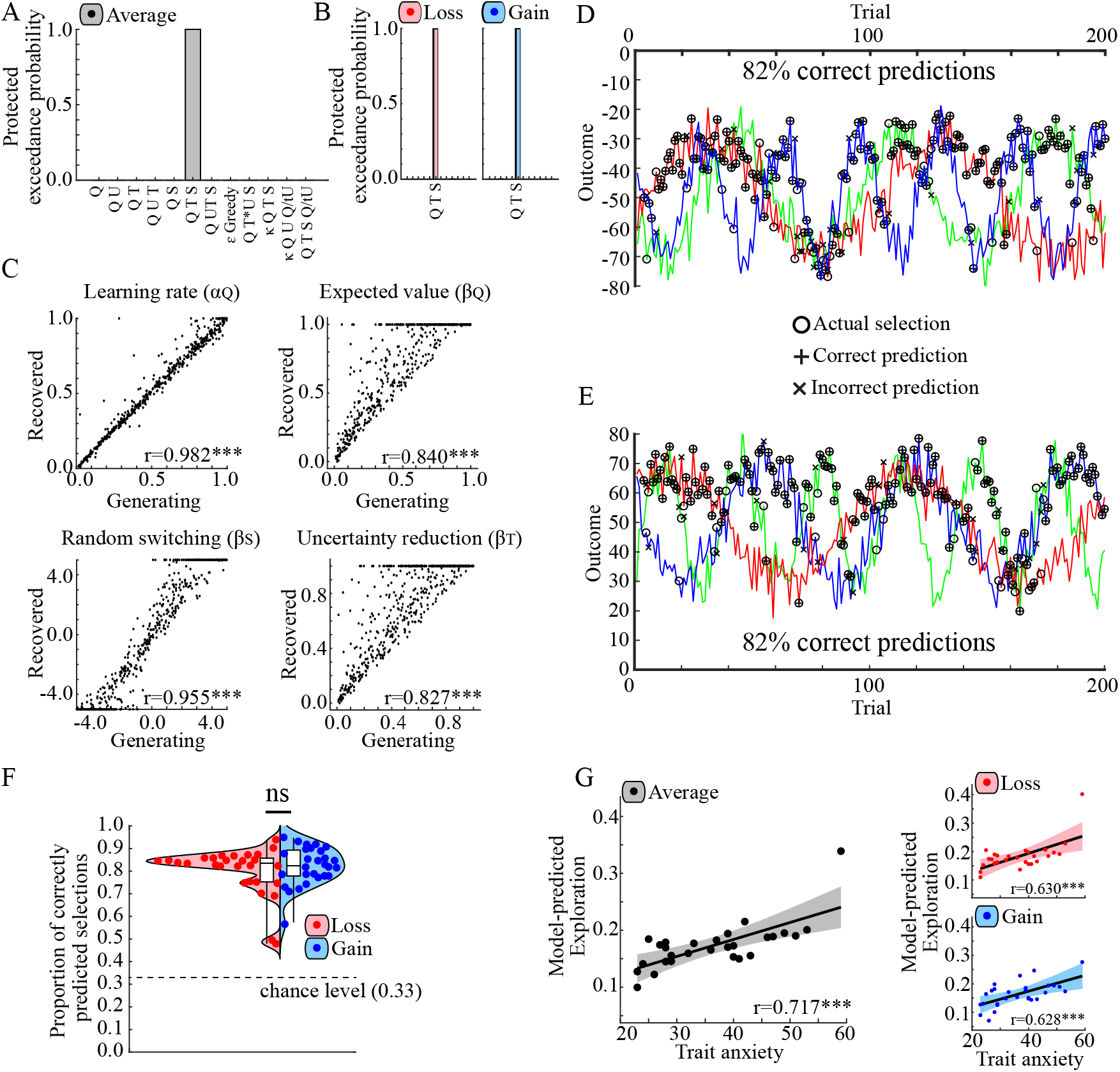
Modeling behavior. **A.** Protected exceedance probabilities obtained from the Bayesian Model Selection (BMS) procedure indicates the QTS model as the winning model when model evidence (here, AICc) was collapsed across conditions (top panel), and **B**. separately in the Loss (left panel) and in the Gain (right panel condition). **C.** The parameter-recovery procedure for the QTS model indicates significant correlations between the parameters used to generate behavior and the recovered parameters obtained from re-fitting the model to the generated behavior. **D,E.** Representative examples displaying the similarity between actual and model-predicted selections in the Loss (**D**) and in the Gain condition (**E**). **F.** The average proportion of correctly predicted selections by the QTS model was well above chance for all participants. **G.** The average model-predicted probability of exploring correlated positively with trait anxiety across conditions (left panel), and in the Loss (top right panel) and in the Gain condition (bottom right panel). ***p<0.001; *p<0.05; ns p>0.05.

As an additional validation, individually fitted parameters were used to calculate the utility of each machine in each trial and the machine with the highest utility was defined as the model’s prediction. This revealed high similarity between actual and model-based selections (Fig.3D-F). Across participants, the model predicted actual selections well above chance-level (i.e. 0.333) in both the Loss (mean correct±SEM: 0.801±0.02) and the Gain condition (mean correct±SEM: 0.822±0.016).

Because the model was deliberately not fitted to exploratory behaviors per-se (to maintain independence), we tested that the model captured relevant aspects of exploration (Wilson and Collins 2019; Palminteri, Wyart, and Koechlin 2017), by calculating a model-derived probability of switching to a suboptimal machine across all trials and conditions. Reproducing the behavioral results, this model-derived probability of exploration was modulated by anxiety (repeated measures two-way ANOVA, main effect of anxiety as linear covariate, Fig.3G, left panel; F(1,26)=27.52, p<0.001, 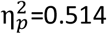, ANOVA and no valence*anxiety interaction effect, F(1,26)=0.16, p=0.696, 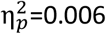, ANOVA). Moreover, the model-derived probability of exploration correlated positively with anxiety in both the Loss (Fig.3G, top right panel; r=0.630, p<0.001, two-tailed Pearson correlation) and the Gain condition (Fig.3G, bottom right panel; r=0.628, p<0.001, two-tailed Pearson correlation). These results mirror the aforementioned behavioral results (Fig.2A).

### Exploration in anxiety trades off value for uncertainty reduction

We tested how the different factors contribute to exploratory decisions and their modulation by anxiety. We repeated the analysis separately for each of the four different factors (namely, for their contribution as quantified by β_*Q*_, β_*T*_, β_*S*_, and α_Q_; Supplementary Note 6). All four factors were significant predictors of behavior (Fig.4A; all p-values for their respective ANOVA intercept terms<0.001; Supplementary Table 3), but only two of them correlated with individual anxiety levels.

**Figure 4.**
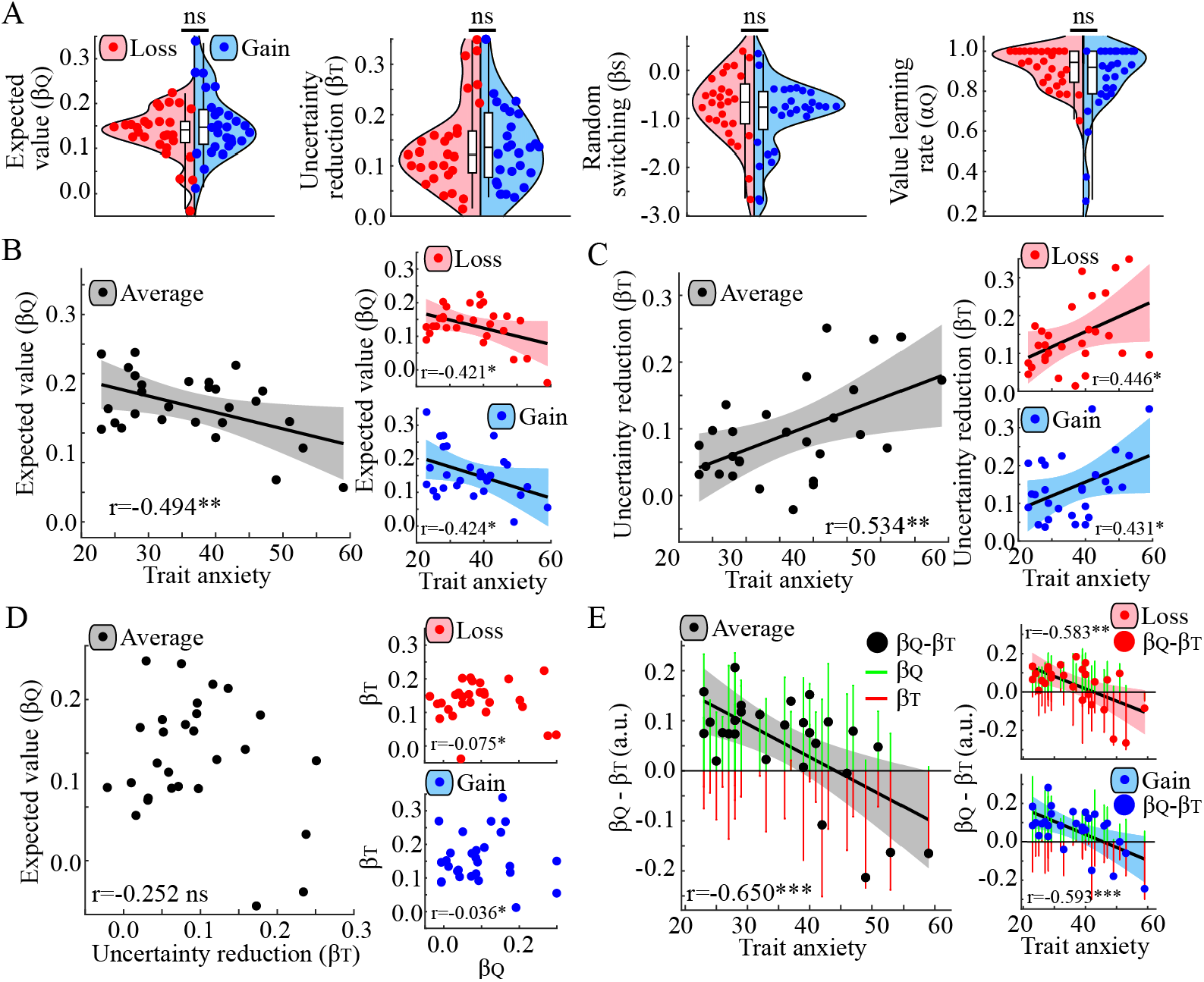
Individually fitted model parameters. **A.** Each of the four fitted parameters of the QTS model are relevant predictors of behavior across participants. **B.** Decision weights estimating the influence of expected value (*β*_*Q*_) were negatively correlated with trait anxiety across conditions (left panel), as well as separately in the Loss (top right panel) and in the Gain condition (bottom right panel). **C.** Decision weights estimating reduction of outcome uncertainty (*β*_*T*_) was positively correlated with trait anxiety across conditions (left panel), as well as separately in the Loss (top right panel) and the Gain condition (bottom right panel). **D.** *β*_*Q*_ did not correlate with *β*_*T*_ across conditions (left panel), nor in the Loss (top right panel) or in the Gain condition (bottom right panel). **E.** The difference between *β*_*Q*_ and *β*_*T*_ correlated negatively with trait anxiety across conditions (left panel), as well as separately in the Loss (top right panel) and in the Gain condition (bottom right panel). ***p<0.001; **p<0.01; *p<0.05; ns p>0.05.

First, expected-value (β_*Q*_) was significantly modulated by anxiety (Fig.4B, left panel; F(1,26)=8.41, p=0.008, 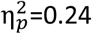, ANOVA; Supplementary Table 3). Specifically, anxiety was negatively correlated with expected-value in both the Loss (Fig.4B, top right panel; r=−0.421, p=0.026, two-tailed Pearson-correlation) and the Gain condition (Fig.4B, bottom right panel; r=−0.424, p=0.025, two-tailed Pearson-correlation; Supplementary Table 3). Additionally, expected-value was negatively correlated with the proportion of exploratory decisions in both the Loss (Supplementary Fig.7A, left panel; r=−0.704, p<0.001, two-tailed Pearson-correlation) and the Gain condition (Supplementary Fig.7A, right panel; r=−0.585, p=0.005, two-tailed Pearson-correlation). This implies that the expected value of different options has a less pronounced impact on decisions made by more anxious individuals.

Second, outcome-uncertainty (β_*T*_) was also significantly modulated by anxiety (Fig4C, left panel; F(1,26)=10.36, p=0.003, 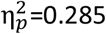, ANOVA; Supplementary Table 3), but in contrast to the negative relationship observed between anxiety and expected-value, outcome-uncertainty was positively correlated in both the Loss (Fig.4C, top right panel; r=0.446, p=0.018, two-tailed Pearson correlation) and the Gain condition (Fig.4C, bottom right panel; r=0.431, p=0.022, two-tailed Pearson correlation). Similarly, outcome-uncertainty correlated positively with the proportion of exploratory decisions in both Gain and Loss (Supplementary Fig.7B; Loss: r=0.412, p=0.029; Gain: r=0.589, p<0.001, two-tailed Pearson correlations). These results imply that more anxious individuals showed a stronger tendency to reduce uncertainty.

Notably, outcome-uncertainty was not correlated with expected-value (Fig.4D; Collapsed: r=−0.252, p=0.19; Loss: r=−0.075, p=0.71; Gain: r=−0.035, p=0.86, two-tailed Pearson correlation), but anxiety was significantly correlated with the difference between these two factors (namely, β_*Q*_-β_*T*_), across condition (Fig.4E, left panel; r=−0.650, p=0.001, two-tailed Pearson correlation), and separately in both the Loss (Fig.4E, top right panel; r=− 0.583, p=0.001, two-tailed Pearson correlation) and the Gain condition (Fig.4E, bottom right panel; r=−0.593, p<0.001, two-tailed Pearson correlation).).

Finally, anxiety did not correlate with random-switching (β_*S*_, Supplementary Table 3; Supplementary Fig.6A,B), or with the learning rate (α_Q_, Supplementary Table 3; Supplementary Fig.6C,D), and neither factors (β_*S*_ or α_Q_) correlated with the proportion of exploratory decisions (Supplementary Fig.7C,D).

To further assess the robustness of these results, we tested the fitted model-parameters of a Kalman-model with a dynamic learning rate, but with the same decision weights as the QTS model (i.e. β_*Q*_, β_*T*_, and β_*S*_; see Methods). Although this model showed an inferior fit to behavior compared to the QTS model (Fig.3A; Supplementary Note 5), the main results were replicated revealing positive correlations between anxiety and the proportion of exploratory decisions, and positive correlations between trait anxiety and β_*T*_ (but not with β_*Q*_; for a summary of these results, see Supplementary Note 7).

Taken together, the results suggest that increased levels of anxiety enhance exploration by trading off immediate expected-value gains for reduction in uncertainty.

### Encoding of prediction-error is not modulated by anxiety

We first tested whether anxiety modulated the neuronal correlates of value learning. As expected from multiple previous studies, activity in the bilateral NAcc correlated significantly with prediction-errors (*δ*_*Q*_; Supplementary Note 8; left NAcc: peak voxel MNI xyz = −9 11 −5, *t*(27)=5.468, p<0.001; FWE, SVC; right NAcc: peak voxel MNI xyz = 9 11 −5, *t*(27)=6.013, p<0.001; FWE, SVC). However, neither anxiety nor valence (gain/loss) impacted the neuronal correlate of predictions-errors (β_δ_ for NAcc ROIs, ANOVAs with within-subject factor valence and anxiety as continuous covariate; Supplementary Note 8). Importantly, these brain activations, in combination with the non-significant correlations between anxiety and learning rates (Supplementary Note 6), suggest that trait anxiety did not affect learning performance in the present task.

### Functional localization of brain regions involved in exploration

To examine the neural representation of expected-value and outcome-uncertainty, we first derived functional ROIs by independently localizing brain regions engaged by exploration-exploitation decisions on a group-level (Vul et al. 2009). In accordance with previous studies (Blanchard and Gershman 2018; Daw et al. 2006; Laureiro-Martinez et al. 2015), exploration was associated with increased activity in the dACC (Fig.5A,B; peak voxel MNI xyz = 9 20 45, *t*(27)=9.717, p<0.001, one-tailed *t*-test FWE SVC), the bilateral aINS (Fig.5C-E; Left aINS: peak voxel MNI xyz = −30 23 3, *t*(27)=8.247, p<0.001; Right aINS: peak voxel MNI xyz = 33 23 3, *t*(27)=9.099, p<0.001, one-tailed *t*-test FWE SVC), and the right FPC (Fig.5F,G; peak voxel MNI xyz = 27 53 10, *t*(27)=5.724, p=0.004, one-tailed *t*-test FWE SVC). On the other hand, exploitation was associated with increased activity in the vmPFC (Fig.5H-I; peak voxel MNI xyz = 0 44 −15, *t*(27)=6.452, p<0.001, one-tailed *t*-test FWE SVC).

**Figure 5.**
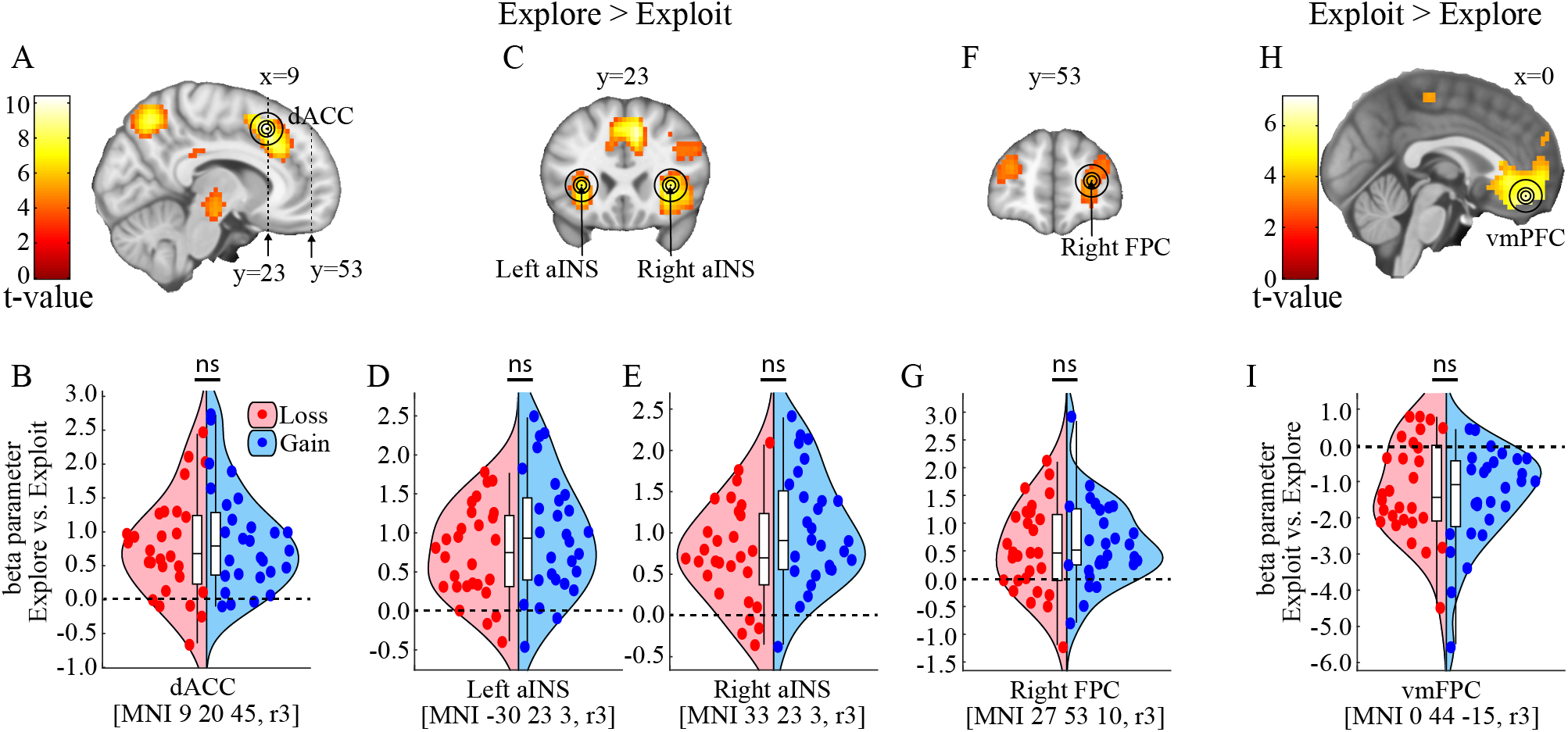
BOLD signal as a function of exploratory decision making. **A-G.** Decisions to explore (vs. exploit) increased activity in (**A,B)** the dorsal anterior cingulate cortex (dACC), (**C-E**) bilaterally in the anterior insula (aINS), and (**F,G**) in the right frontopolar cortex (FPC). Extracted beta parameters for decisions to explore (vs. exploit), extracted from spheres centered on the peak activation within a priori ROIs in the dACC (**B**), L aINS (**D**), R aINS (**E**), and the R FPC (**G**). **H.** Decisions to exploit (vs. explore) increased activity in the ventromedial prefrontal cortex (vmPFC). **I.** Extracted beta parameters for decisions to exploit (vs. explore), extracted from 3 mm radius spheres centered on the peak activation within the vmPFC ROI. Peak voxel coordinates presented here were used in subsequent analyses by extracting the average voxel intensity within 3 mm spheres centered on these coordinates. To ensure the robustness of any observed effects, all analyses were repeated using spheres with different radii (0, 3, 5, and 10mm), as illustrated in **A**, **C**, **F**, and **H**. For visualization purposes, the brain activation in **A**, **C**, **F**, and **H**, is displayed using an uncorrected threshold of p=0.001. ns p>0.05.

We then used individual trial-by-trial differences in expected-value (*ΔQ*) and outcome-uncertainty (*ΔT*), between the selected machine and the average of the two rejected machines, as parametric modulators to identify their respective brain correlates (*β*_Δ*Q*_, *β*_Δ*T*_). Activity estimates were extracted from 3mm radius spheres centered on the peak voxels identified above (for robustness and validation, analyses were repeated using spheres with different radii of 0, 3, 5, and 10mm, all producing the same results described below, see Methods), and were used in separate repeated measures ANOVAs with within-subject factor valence and anxiety as a continuous covariate.

### Anxiety modulates the representation of expected-value in the dACC

In the dACC, anxiety correlated significantly with the representation of expected-value *β*_Δ*Q*_ (Fig.6B, left panel F(1,26)=10.242, p=0.004, 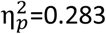, ANOVA), but not in any other region (Supplementary Note 9). Specifically, anxiety correlated negatively with *β*_Δ*Q*_ in both the Loss (Fig.6B, top right panel; r=0.375, p=0.049, two-tailed Pearson correlation) and in the Gain condition (Fig.6B, bottom right panel; r=−0.603, p<0.001, two-tailed Pearson correlation). Across participants, the average value of *β*_Δ*Q*_ was positive (F(1,26)=60.594, p<0.001, ANOVA intercept term; Supplementary Table 5). Because *β*_Δ*Q*_ reflects the coupling between BOLD signal in the dACC and expected value differences *ΔQ* (Fig.6C), these results mean that participants with a large *β*_Δ*Q*_ (e.g. low anxiety) show an increased engagement of the dACC when *ΔQ* decreases, while the dACC of more anxious individuals is less responsive to changes in *ΔQ* (*β*_Δ*Q*_ close to zero).

**Figure 6.**
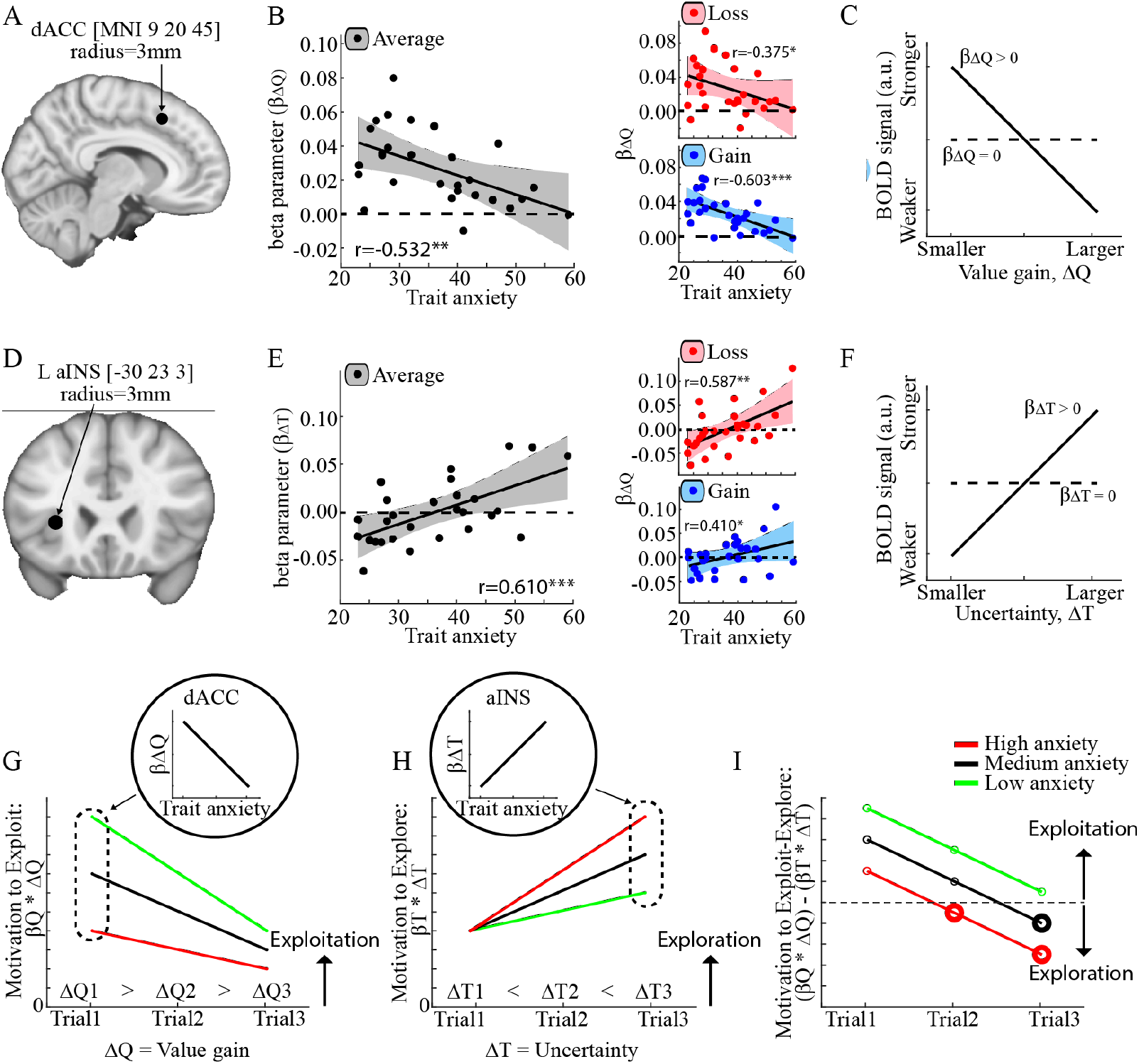
**A-F.** Brain correlates of differences in expected value (i.e. *ΔQ*) and uncertainty (i.e. *ΔT*) that is significantly modulated by Trait anxiety. Beta parameters were extracted from 3mm radius spheres (indicated by black dots in **A** and **D**) centered on the coordinates obtained from the independent localizer. **B.** *β*_Δ*Q*_ correlated negatively with trait anxiety in the dACC across conditions (left panel), and separately in the Loss (top right panel) and in the Gain condition (bottom right panel). **C.** To understand these results, consider that a positive *β*_Δ*Q*_ indicates that dACC activity increases when the expected values becomes increasingly similar, while a *β*_Δ*Q*_ close to zero (e.g. high anxiety) means that dACC activity does not change with *ΔQ*. **E.** *β*_Δ*T*_ correlated positively with trait anxiety in the bilateral aINS (here, only the left aINS is shown) across conditions (left panel), and separately in the Loss (top right panel) and in the Gain condition (bottom right panel). **F.** To understand these results, consider that a positive *β*_Δ*Q*_ (e.g. high anxiety) indicates that aINS activity increases when the uncertainty increases, while a *β*_Δ*Q*_ close to zero means that aINS activity does not change with *ΔT*. **G-I.** Schematic summary of the results. Consider a toy-example where the values of the machines are initially known (e.g. 40, 15, 25) and the best machine is selected in three consecutive trials which yield decreasing outcomes (e.g. 35-30-25). Accordingly, the differences in expected value (*ΔQ*) gradually decrease (**G**) while uncertainty of the non-selected machines (*ΔT*) increase (**H**). **G.** The motivation to exploit the best machine in a given trial is estimated by *ΔQ* scaled by an individual’s weighting of expected-value gains (*β*_*Q*_), i.e. *ΔQ* ∗ *β*_*Q*_, where *β*_*Q*_ is smaller in high anxiety. The weighting of expected values may relate to the representation of ΔQ in the dACC, i.e. *β*_*ΔQ*′_, which was close to zero in anxious individuals. **H.**The motivation to explore is estimated by *ΔT* scaled by an individual’s weighting of uncertainty reduction (*β*_*T*_), which was larger in high anxiety, i.e. *ΔT* ∗ *β*_*Q*_. The weighting of uncertainty may relate to the representation of ΔT in the aINS, i.e. *β*_*ΔT*_, which was larger in more anxious individuals. **I.** The exploit-explore trade-off is determined by the difference in the motivation to exploit versus explore, which is shifted towards exploration for more anxious individuals. ***p<0.001; **p<0.01; *p<0.05; ns p>0.05.

We tested if the reduced tracking of expected-value differences in the dACC can be attributed to an overall increase in dACC activity, which could indicate a limited ability to encode task-related parameters due to dACC saturation. However, overall dACC activation did not correlate with anxiety, suggesting against this possibility (Supplementary Note 10). Additionally, anxiety did not affect the representation of expected-value in the vmPFC (Supplementary Note 9), a brain region directly involved in value representations (Levy and Glimcher 2012; Domenech and Koechlin 2015).

Overall, anxiety modulates the neuronal representation of expected-value differences in the dACC. This result parallels and extends the behavioral finding that decisions made by more anxious individuals were less influenced by the immediate expected-value.

### Anxiety modulates the representation of outcome-uncertainty in the aINS

In the aINS, anxiety correlated significantly with the representation of outcome-uncertainty difference *β*_Δ*T*_ (Fig.6E, left panel; left aINS: F(1,26)=15.432, p<0.001, 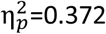, ANOVA; right aINS: F(1,26)=11.855, p=0.002, 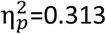, ANOVA), but not in any other region (Supplementary Note 11). Here, anxiety was positively correlated with the representation of outcome-uncertainty in the aINS in both the Loss (Fig.6E, top right panel; left aINS: r=−0.587, p=0.001; right aINS: r=−0.492, p=0.008, two-tailed Pearson correlations) and the Gain condition (Fig.6E, bottom right panel; left aINS: r=−0.410, p=0.030; right aINS: r=−0.345, p=0.028, two-tailed Pearson correlation). To understand this result, consider that a positive value of *β*_Δ*T*_ indicates increased BOLD signal for an increased uncertainty *ΔT* (Fig.6F). Therefore, the aINS of more anxious individuals (large positive *β*_Δ*Q*_) is more responsive to increases in uncertainty.

Overall, anxiety modulates the neuronal representation of outcome-uncertainty differences in the bilateral aINS. This result parallels and extends the behavioral finding that decisions made by more anxious individuals were more influenced by outcome-uncertainty.

### The neuronal trade-off mediates the behavioral trade-off

Finally, we tested whether the behavioral tradeoff between expected-value and uncertainty-reduction observed in more anxious individuals (i.e. *β*_*Q*_ − *β*_*T*_; Fig.4E) is related to a differential representation of expected-value in the dACC (Fig.6B) and outcome-uncertainty in the aINS (Fig.6D). We found that the strength of the relationship between anxiety and the difference between expected-value and uncertainty-reduction, decreased significantly when controlling for the differential representation in the two brain regions. This was the case in both the Loss and the Gain condition (Loss: Δb_Anxiety_=−0.254, p<0.001, Gain: Δb_Anxiety_=−0.292, p<0.001; Mediation analysis with Monte-Carlo randomization test; Supplementary Note 12).

Overall, these results indicate that the behavioral tradeoff between expected-value and uncertainty-reduction, which determines the anxiety-related exploration rate, is mediated by how the brain represents expected-value in the dACC versus outcome-uncertainty in the aINS.

## Discussion

We show here that individuals with higher levels of trait anxiety tend to explore more in a dynamic decision-making environment, and do so in a similar manner in both Gain and Loss conditions. This exploration-exploitation bias was associated with two distinct neuro-behavioral mechanisms. First, more anxious individuals were less influenced by immediate expected-values and showed a reduced correlation between activity in the dACC and expected-value differences. Second, more anxious individuals were more influenced by outcome-uncertainty and showed an increased correlation between outcome-uncertainty and activity in the aINS (see Fig.6G-I for a schematic summary of the results). Taken together, anxiety boosts exploration by trading off immediate value gains in order to reduce the uncertainty of future decisions, paralleled by a tradeoff in dACC-aINS activity.

Our results provide evidence that individual anxiety levels contribute to the trade-off between immediate benefits, namely gaining a reward or avoiding a risky/uncertain situation, and prospective decision making. In the current study, as in most real-life situations, exploration provides information that benefits future decision making. Our findings therefore extend on previous studies that found that anxiety increases aversion for uncertainty, both in terms of risk (i.e. known uncertainty) and ambiguity (i.e. unknown uncertainty; (Grupe and Nitschke 2013; Hartley and Phelps 2012). In these previous studies, however, there was no information to be gained from making a risky decision (e.g. (Charpentier et al. 2017)). Taken together, the evidence suggests that whereas anxiety decreases risk-taking in an environment where all information is provided and nothing can be learned, anxiety increases the willingness to confront uncertainty via increased exploration in a dynamic environment, where risky decisions provide prospective benefits in the form of information gains and/or uncertainty reductions.

Here, anxious individuals were more sensitive to a particular type of uncertainty, one that depends on the dynamics of the environment, and which is not present in static environments (e.g. where reward contingencies do not change across trials). Related to this finding, it has been shown that exploration depends on the overall uncertainty of the environment, such that exploration increases in more volatile and uncertain settings (Knox et al. 2011; Gershman 2018; Gershman and Tzovaras 2018). Combined with the present results, one possible hypothesis is that anxious individuals perceive the world as being more volatile and therefore needs to be explored more frequently. Previous studies reported that trait anxiety and internalizing psychopathology decreased the ability to adjust learning rates as a function of volatility (Browning et al. 2015; Gagne et al. 2020), and it has been suggested that anxious individuals misestimates uncertainty (Pulcu and Browning 2019). Indeed, one simple heuristic which can be used to maintain a low level of uncertainty in a decision-making environment is to explore frequently, independently of actual volatility levels.

Of note, neither outcome-variability nor total outcome-variability contributed to exploration in the present study. This finding is consistent with results obtained in previous studies using similar ‘restless n-armed bandit tasks’, e.g. dynamic decision making tasks (Daw et al. 2006; Payzan-LeNestour and Bossaerts 2011), but different from findings in other tasks where reward-contingencies were static and outcome-variability was suggested to drive directed exploration, such as in the ‘clock-task’ (Badre et al. 2012; Frank et al. 2009), or specific instances of two-armed bandit-tasks in which direct and random exploration was related to outcome-variability and total outcome-variability (Tomov et al. 2020; Gershman 2018; Gershman and Tzovaras 2018). One reason for the differences could be that dynamic environments require the continuous monitoring and updating of values, and their respective uncertainties, across all trials of an experiment, and therefore entails a heavier working memory load. By contrast, in static environments most of the learning occurs within the first few trials. Indeed, reinforcement learning has been shown to interact with working memory, such that the ability to make value-based decisions depends on working memory load (Collins and Frank 2012). As such, the reduced influence of outcome-variability on decision making in dynamic environments could be either due to a degraded/noisier representation of uncertainty in the working memory, and/or the application of simpler decision-making strategies that reduces working memory load, such as keeping track of only expected-value but not outcome-variability.

While directed and random exploration was not related to outcome-variability in the present study, high trait anxiety increased exploration via more uncertainty-reduction and by decreasing the weighting of expected-value. An alternative interpretation of the present results therefore, framed in the context of directed and random exploration, is that increased uncertainty-reduction can be regarded as a form of ‘directed’ exploration because it reduces the uncertainty of a particular option, while a reduced focus on expected-value could be seen as a form of ‘random’ exploration because it allows deviations from purely greedy decisions. In other words, high trait anxiety may increase both directed and random exploration in dynamic environments.

The positive correlations between uncertainty-reduction and anxiety are in accordance with the previously described link between anxiety and intolerance to uncertainty (Buhr and Dugas 2009; Dar, Iqbal, and Mushtaq 2017), and the idea that anxiety is underpinned by the fear of the unknown (Carleton 2016). Yet, whether the fear of the unknown is relevant only in contexts with aversive outcomes (as opposed to also in positive contexts) is currently under debate (Einstein 2014; Grupe and Nitschke 2013). Our results, showing equal contributions from uncertainty in Gain and Loss conditions, suggest that it is the uncertainty itself, and not its interaction with outcome valence, that promotes exploration. Corroborating this notion, a recent study exposed participants to different types of written real-life scenarios with uncertain positive or negative outcomes, and revealed similar correlations between trait anxiety and aversion to uncertainty in both scenarios (Pepperdine, Lomax, and Freeston 2018), further supporting the notion that anxiety is associated with a general aversion to uncertainty. An interesting extension of these findings could be that changing an individual’s intolerance to uncertainty in a positive context, for example via cognitive bias modification (Oglesby, Allan, and Schmidt 2017), may cause a general reduction in intolerance to uncertainty that transfers to negative contexts.

At the neuronal level, trait anxiety correlated negatively with the representation of differences in expected-value in the dACC, yet this activity increased overall across participants when expected-values became increasingly similar. This latter result is in accordance with findings obtained in foraging paradigms, where the dACC was found to track the relative difference in expected-value between continued exploitation and foraging (defined as a decision to abandon gradually depleting options; (Kolling et al. 2012; Shenhav et al. 2014)). Foraging theory posits that a decision to forage is optimal when the expected value difference between an exploited option and foraging options is small (Hayden, Pearson, and Platt 2011; Charnov 1976). Likewise, exploration is arguably optimal when the expected values of options are similar, because at this point additional information is needed to deduce which is the best option. A more accurate representation of relative expected-values in the dACC should therefore bias behavior towards exploitation; and correspondingly, a less accurate representation as seen in anxious individuals, should bias behavior towards exploration. Notably, it was recently argued that non-greedy decisions could result from noisy learning processes, where the amount of learning noise was tracked by the dACC well into the decision process (Findling et al. 2019). However, if decisions made by more anxious individuals in the present study was affected by such learning noise, we would expect to see an overall increase in dACC activation as a function of trait anxiety during decision making, but this was not the case. Moreover, we included an explicit factor in our model, outcome-uncertainty, which was uncorrelated with expected-value, and which significantly contributed to explain the proportion of non-greedy, exploratory decisions elicited by more anxious individuals.

Furthermore, anxious individuals showed a more positive coupling between outcome-uncertainty and BOLD signal in the bilateral aINS, as well as increased uncertainty reduction. These results parallels findings showing that the aINS responds to different types of uncertainty (Simmons et al. 2008), is involved in the aversion and the intolerance of uncertainty (Grupe and Nitschke 2013), and is hyper-responsive to uncertainty and distal threats in high anxiety (Etkin and Wager 2007; Grupe and Nitschke 2013; Fung et al. 2019b). Accordingly, a more rapid accumulation of uncertainty-related activation in the aINS facilitates decisions to explore, as displayed by more anxious individuals. Because the aINS also represents negative affective states and stimuli, including pain, disgust, anxiety, conditioned fear, negative pictures, and odors (Gogolla 2017; Alvarez et al. 2015; Ploghaus et al. 1999; Wright et al. 2004), a putative mechanism by which the aINS can increase exploration might be that uncertainty-related activation of the aINS is perceived as aversive, leading to decisions aimed at reducing uncertainty, namely exploration, and hence avoiding the negative state. This hypothesis needs to be tested directly.

These ideas are similar to recent findings in studies of over-generalization (Dunsmoor and Paz 2015; Lissek 2012). There as well, aversive contexts led to wider stimulus generalization (Laufer and Paz 2012; Resnik, Sobel, and Paz 2011), similar to our finding here of an overall higher exploration in aversive contexts; yet in anxious individual, these studies found wider generalization for both aversive and appetitive contexts (Laufer, Israeli, and Paz 2016), similar to what we find here that exploration is increased with anxiety for both types of valence. Because wider/over-generalization can also stem from uncertainty and less information about the stimulus, the two mechanisms might be linked. One possibility is an attentional-shift between top-down processes devoted to value-based decision making - evidenced by weaker dACC activity, to bottom-up processing of aversive information of increased uncertainty - evidenced by a responsive aINS (Eysenck et al. 2007; Bar-Haim et al. 2007). Supporting this idea, we found that the increased trade-off between expected-value and uncertainty-reduction, as seen in individuals with high trait anxiety, was mediated by the differential neuronal representation of expected-value in the dACC and uncertainty-reduction in the aINS.

Exploratory decision making is an integral part of everyday life, being involved in decisions ranging from small (e.g. where to eat lunch) to large (e.g. which partner to marry). Therefore, our results highlight two mechanisms which may be critical determinants of life-quality in anxious individuals, even if in the normal-range. A high exploratory drive could reduce the ability to engage in long-term commitments, and a reduced focus on expected values may lead to faster abandoning of good positions. In turn, maladaptive exploratory strategies could trigger a cascade of events that reinforces symptoms of anxiety and depression. For example, exaggerated exploration may lead to increased exposure to suboptimal outcomes which could trigger feelings of regret (Coricelli and Rustichini 2010) and reduce confidence in one’s ability to make appropriate decisions (Bandura 1982). Indeed, anxiety and depression have been associated with both increased feelings of regret (Roese et al. 2009; Ehring and Watkins 2008) and reduced self-efficacy (Muris 2002; Zumberg, Chang, and Sanna 2008; Luszczynska, Gutierrez-Dona, and Schwarzer 2005). Moreover, high levels of trait anxiety is a risk-factor for chronic anxiety and depression (Chambers, Power, and Durham 2004; Weger and Sandi 2018), and their development and maintenance may depend on dysfunctional learning and decision making (Mineka and Oehlberg 2008; Paulus and Yu 2012; Lee 2013; Rosen and Schulkin 1998; Grupe 2017).

We estimated anxiety using the standard Spielberger’s Trait-Anxiety Inventory (Spielberger et al. 1983), mainly because it provides a gradual scale also for the normal (below clinical) range. This provides two benefits: first, observing how exploration varies across the distribution, rather than comparing two populations which are somewhat arbitrarily divided (patients vs. controls); and second, examining the normal range, allowing us to examine how maladaptive decisions are mediated by exploration even in healthy individuals. Because such maladaptive decision have a huge impact on daily-life in all individuals, and definitely on societies and industry, we argue that more studies should use such gradual scales over non-clinical populations (Browning et al. 2015; Fung et al. 2019a; Gagne et al. 2020). It should be noted that Spielberger’s Trait-Anxiety Inventory has been debated for its lack of convergent and discriminant validity, suggesting that it estimates ‘negative affectivity’ rather than proneness to anxiety per-se (Balsamo et al. 2013). Yet, because negative affectivity is closely linked to psychopathology (Kotov et al. 2010; Stanton and Watson 2014), and has been noted as a vulnerability factor for developing anxiety and depression (Clark, Watson, and Mineka 1994), our results still bear significant relevance.

In summary, we identify two neuro-behavioral mechanisms whose tradeoff determines exploratory bias and is directly related to anxiety levels. Importantly, we show that anxiety within the ‘normal’ spectrum is a critical determinant of exploratory behaviors in humans, might increase maladaptive decision-making, and therefore affect life-quality in individuals dealing with everyday decisions. We suggest that in some individuals, these factors can develop further into a vicious cycle. Put simply, poor decision making caused by an excessive drive to explore may increase the frequency of poor real-life outcomes, which in turn may contribute to psychopathology.

## Acknowledgments

We thank Dr. Edna Furman-Haran and Fanny Attar for MRI procedures. K.C. Aberg was supported by the Swiss Society of Friends of the Weizmann Institute Postdoctoral Fellowship grant. The work was supported by a Joy-Ventures grant, an ISF #2352/19 and an ERC-2016-CoG #724910 grants to R. Paz.

## Financial Disclosures section

The authors declare no competing financial interest.

## Methods

### Participants

After having provided written informed consent, fifty-one (51) participants (n=20/31 for the behavioural pilot/main-fMRI study) joined the experiment. The study was approved by the Helsinki committee of the Sourasky Medical Center, under protocol number 0287–09-TLV (Ministry of health protocol no. HT5271) and further approved by the Institutional Review Board (IRB) of the Weizmann Institute.

All participants were right-handed and reported no previous history of psychiatric or neurological disorders. Handedness was assessed by asking participants to indicate their preferred hand for writing and drawing, on a scale ranging from 1 (always using the left hand) to 7 (always using the right hand). Only participants that indicated always using the right hand on both questions were recruited. Two participants who fell asleep in the scanner, and one participant who displayed excessive head movements during scanning were excluded. Therefore, data from 28 participants (eight males; average age ± SEM: 27.571 ± 0.730) were included in the subsequent data analyses. These participants showed a normative distribution of trait-anxiety scores (mean=36.321, SD=9.982; see (Spielberger et al. 1983), Table 1).

### Task

In each trial, participants decided which of three slot machines to play (Fig.1A,B). The outcome of the selected machine was then revealed. Exploration was promoted by varying the outcome assigned to each slot machine across trials (Fig.1C). Thus, to optimize overall performance participants needed to track the expected outcome associated with each slot machine, and to make exploratory decisions to detect outcome changes.

Each trial started with a fixation cross displayed for 500 ms, followed by the display of the three slot machines (Fig.1A,B). Participants needed to elicit a response within 1500 ms, after which the selected machine became animated. The animation lasted for 1500 ms minus the response time plus a jittered time duration drawn from an exponential distribution with mean 3500 ms. The outcome associated with the selected machine was then presented for 1000 ms (Fig.1A,B), after which the next trial was initiated following a jittered delay drawn from an exponential distribution with mean 3500 ms. If no response had been registered within the 1500 ms, the words ‘Too slow’ were displayed for 1000 ms.

The outcome distribution of each slot machine was described by different sine-wave functions (some examples are shown in Fig.1C-G). The magnitude and mean of each sine-wave was always 20 and 50 points respectively. The period lengths of the three sine-waves in a session were 100, 67, or 33 trials, respectively, and the sine-waves were phase-shifted by 0.10, 0.45, and 0.90 period lengths. A pseudorandomization scheme was administered to ensure that no participant was exposed to the same stimulus configuration by assigning different phase-shifts to each sine-wave, different sine-waves based on Gain/Loss conditions, spatial position, and color. Moreover, the assignment of color to a machine’s spatial position and Gain/Loss condition also varied pseudorandomly.

Participants engage more in risky decisions when facing certain losses, as compared to certain gains (De Martino et al. 2006), and this effect is exaggerated in more anxious individuals (Jepma and Lopez-Sola 2014; Xu et al. 2013). Here, to test for differences in exploratory decision making between gain and loss conditions, each participant performed two versions of the task. In the Gain condition, all outcomes were positive and participants acquired points (Fig.1D,F). By contrast, participants had to avoid losing points in a Loss condition with only negative outcomes (Fig.1E,G). The points corresponded to a monetary bonus at the end of the experiment. In the Gain condition, the bonus (initially set to 0 Israeli new shekels) increased as more points were collected. In the Loss condition, the bonus (initially set to 10 shekels) decreased with the accumulation of negative points. At the end of the experiment, each participant’s bonus was calculated as 10 * proportion of maximal and minimal points for the Gain and Loss condition, respectively. The order of Gain and Loss conditions was counterbalanced across participants.

### Modelling approach

As in previous studies, a computational approach was used to determine whether a decision could be classified as exploratory or as an exploitation (Daw et al. 2006). Specifically, an exploratory decision was defined as the selection of a suboptimal machine which was different from the machine selected in the previous trial (Tzovara et al. 2012), where suboptimal referred to any machine with a less-than-highest expected value.

For all models, the expected value *Q*_*i*_(*t*) of the selected machine *i* in trial *t* was updated by the mismatch between the expected value and the actual outcome *R(t)*, i.e. the reward prediction error δ_*Q*_(*t*), scaled by the learning rate α_*Q*_.

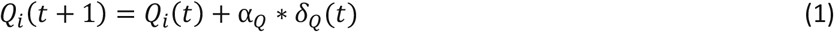

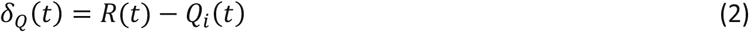

The probability *p* of selecting machine *i* in trial *t* was modeled using two approaches. For all models except one (see the ε-greedy model below), decisions were modeled using a soft-max choice probability function in which the probability of selecting a particular machine depends on its utility, u_*i*_:

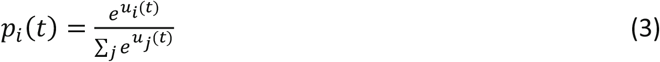

Here, u_*i*_ is defined as the sum of parameters that may contribute to a decision (e.g. expected value) multiplied by their respective decision weights (e.g. how much expected value contributes to a decision). For example, one common way to model exploratory tendencies is by the inverse temperature β_*Q*_. Specifically, the decision weight β_*Q*_ determines the trade-off between exploitation and exploration such that small and large values of β_*Q*_ are associated with increased and decreased exploration, respectively, because β_*Q*_ determines the influence of expected value on a decision:

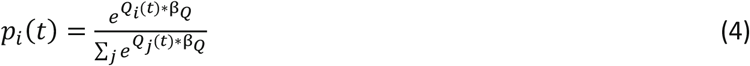

However, other factors unrelated to the expected values of stimuli may also contribute to decision making. For example, exploration may be directed towards options with large outcome-variability (Badre et al. 2012; Frank et al. 2009; Gershman 2018). To test this notion, the outcome-variability *U*_*i*_(*t*) was updated by the mismatch between the expected outcome-variability and the absolute reward prediction error δ_*Q*_(*t*), i.e. the outcome-variability prediction error δ_*U*_(*t*), scaled by the learning rate α_*U*_:

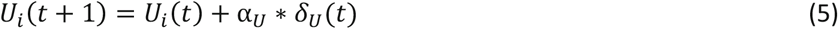

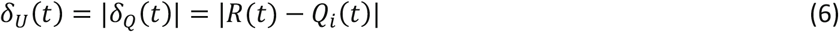

Additionally, in a dynamic environment, the uncertainty associated with an option increases with the time since it was last visited. For example, an outcome is more likely to have changed for a slot machine that has not been selected for many trials. To test the impact of this factor on decision making, we defined outcome-uncertainty *T*_*i*_(*t*) as the number of trials since a machine was last selected.

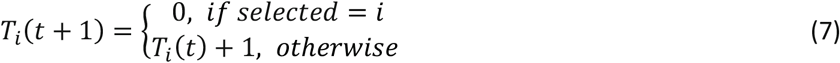

We also modeled the impact of an individual’s tendency to randomly switch between different machines, a decision making strategy which could help to rapidly discover changes in the environment. Accordingly, *S*_*i*_(*t*) is set to 0 and 1 for slot machines that were selected and rejected, respectively. We refer to this factor here as ‘random switching’, whereas previous research denotes the reverse of this factor as ‘choice stickiness’ or ‘perseverance’ (Chakroun et al. 2020; Gershman, Pesaran, and Daw 2009).

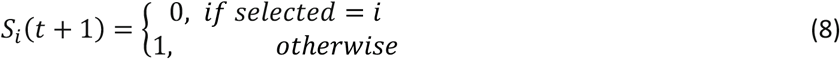

Given reports of more exploration with increased uncertainty in the environment (Gershman 2018; Wilson et al. 2014), we also included a factor which scales the impact of expected value with the total uncertainty of the environment, as estimated by the total outcome-variability (Gershman 2018; Gershman and Tzovaras 2018; Wilson et al. 2014). Specifically, *tU*_*i*_(*t*) is defined as:

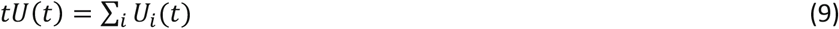

These parameters were included in a soft-max choice probability function. Thus, in the model including all these parameters, the probability of selecting machine *i* depends on β_*Q*_, β_*U*_, β_*T*_, β_*S*_, β_*tU*_, and β_*U*∗*T*_, which determine the influence of expected-value *Q*_*i*_(*t*), outcome-variability *U*_*i*_(*t*), outcome-uncertainty *T*_*i*_(*t*), random-switching *S*_*i*_(*t*), expected-value scaled by total outcome-variability *Q*_*i*_(*t*)/*tU*(*t*), and outcome-uncertainty scaled by outcome-variability *T*_*i*_(*t*) ∗ *U*_*i*_(*t*), respectively.

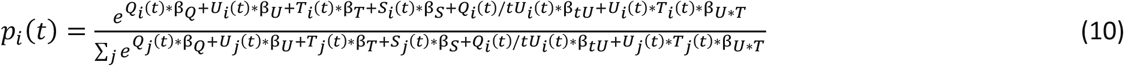

We also implemented an *ε*-greedy algorithm, in which the probability of selecting the machine with the highest expected value in a trial is 1-*ε*. Accordingly, small and large values of *ε* are associated with decreased and increased exploration, respectively:

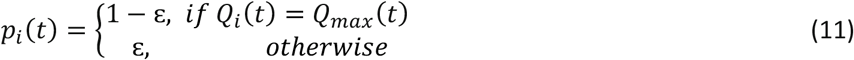

Finally, we included two Kalman-filter models where the learning rate varies between trials as a function of a machine’s current level of uncertainty. Models based on Kalman-filters have been used previously to disentangle random and directed exploration in human decision making (Bishop 2006; Gershman 2018; Gershman and Tzovaras 2018). In these models, the prediction error and expected-value update is determined by Equations 1–2, but the learning rate varies from trial-to-trial according to:

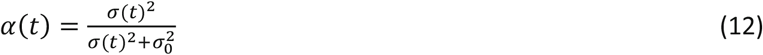

σ(*t*)^2^ and 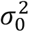 respectively denotes the selected machine’s variance in trial t and prior variance. The variance of the selected machine is updated according to:

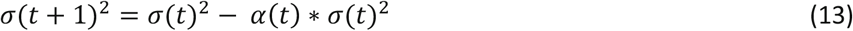

In the first Kalman-filter type model, which is based on previous research (Gershman 2018; Gershman and Tzovaras 2018), decision weights were included for expected-value (β_*Q*_), outcome-variability (β_*U*_; here, the outcome variability is estimated in each trial by σ), and expected-value scaled by total outcome-variability (β_*tU*_; here, tU is estimated in each trial by 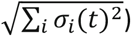:

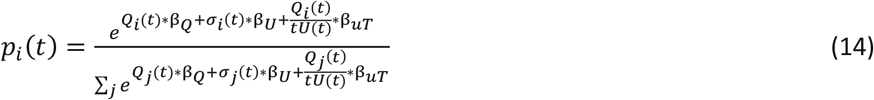

In a second Kalman-filter model, decision weights were included for expected-value (β_*Q*_), outcome-uncertainty (β_*T*_; defined as in Equation 7), and random-switching (β_*S*_; defined as in Equation 8).

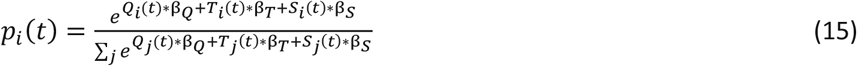

All free parameters were fitted individually to each participant’s behavior by minimizing the negative log-likelihood estimate:

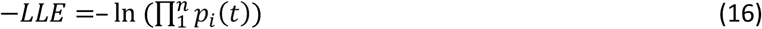

To avoid getting stuck in local minima, each fit was repeated 100 times with different random starting points for each free parameter. The two learning rates were limited by 0 and 1, while no bound was applied for the β decision weights (yet, the random starting points were limited by −5 and +5). For the Kalman-filter models, the prior variance 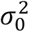 was set to 4.0 for reasons of model-degeneracy (Daw et al. 2006).

To obtain the most parsimonious model, i.e. the model that provides the best trade-off between its complexity and goodness of fit, different models incorporating different combinations of the above mentioned parameters were created and confronted. In total, 12 different models were tested: models ‘Q’, ‘QU’, ‘QT’, ‘QUT’, ‘QS’, ‘QTS’, ‘QUTS’, ‘Q T*U S’, and ‘QTS Q/tU’, all use constant learning rates and differed only in the factors included in the softmax choice-probability function, as indicated by their denotations. ‘κ-QTS’ and ‘κ-QU Q/tU’ are Kalman-filter type models using softmax choice probability function. In other words, the only difference between the QTS and the κ-QTS models is that they use constant and variable learning rates, respectively. Finally, the *ε*-greedy model uses a constant learning rate, but an *ε*-greedy choice probability function (Equation 11).

A version of Akaike’s information criterion (AICc) that penalizes a model’s goodness of fit based on the number of trials *n* and the number of fitted parameters *k* was used to compare the goodness of fit between models.

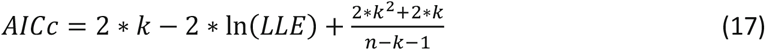

To determine the most parsimonious model based on using the AICc as model evidence, we applied a Bayesian Model Selection (BMS) procedure (Stephan et al., 2009). The BMS assumes that the observed behavior of each participant may have been generated by a different model drawn from an unknown population distribution. The most parsimonious model can then be determined via the protected exceedance probability, which is defined as the posterior probability that the model has a higher frequency than the other tested models, while also accounting for the probability that differences were due to chance. For consistency, we performed the same BMS analysis using the Bayesian Information Criterion (BIC) as model evidence, because the BIC provides a larger penalty for more complex models, i.e. k ∗ ln(*n*) for BIC rather than 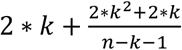 for AICc (Bishop 2006; Gershman 2018).

### Parameter recovery

It needs to be shown that fitted model-parameters are meaningful via their extraction from a data set with known parameters (Wilson and Collins 2019). To accomplish this, the behaviors of 500 virtual participants were simulated using the QTS model by randomly sampling the model parameters from uniform distributions with boundaries [0 1], [0 1], [0 1], and [−5 5] for α_Q_, β_Q_, β_T_, and β_S_.

### Statistical analyses

Behavior and computational results were analyzed using traditional statistical approaches, including repeated measures ANOVAs, *t*-tests, Pearson correlations, and nonlinear regression. A mediation analysis was conducted to test whether the nonlinear relationship between Trait anxiety (T) and Overall performance (O) was mediated by Exploration (E). In essence, this approach tests whether a nonlinear relationship between an independent variable (here, T) and a dependent variable (here, O) is significantly reduced by including a mediating variable (here, E) in the regression analysis (Hayes and Rockwood 2017). Critically, according to contemporary statistical thinking, it is sufficient to show that the predictive value of an independent variable is significantly reduced when controlling for a mediating variable (Hayes and Rockwood 2017). A Monte-Carlo randomization procedure was performed to assess statistical significance: First, the quadratic regression coefficients of T predicting O alone (b^2^_Anxiety1_), and when both T and E predicts O (b^2^_Anxiety2_ and b^2^_Explore_) were calculated. The change in T’s predictive value was calculated as Δb^2^_Anxiety_=b^2^_Anxiety2_-b^2^_Anxiety1_. To assess the statistical significance of Δb^2^_Anxiety_, the above analysis was repeated 10’000 times by each time randomly shuffling the values of E. This procedure creates a distribution (Δb^2^_Anxiety, null_) for the null-hypothesis that Exploration does not change the relationship between Trait anxiety and Overall performance. Statistical significance of the observed Δb^2^_Anxiety_, and therefore also whether Exploration mediated the relationship between Trait anxiety and Overall performance, was assessed by comparing Δb^2^_Anxiety_ to the resulting null-distribution Δb^2^_Anxiety, null_.

### MRI Data

#### Image Acquisition

MRI images were acquired using a 3T whole body MRI scanner (Trio TIM, Siemens, Germany) with a 12-channel head coil. Standard structural images were acquired with a T1 weighted 3D sequence (MPRAGE, Repetition time (TR)/Inversion delay time (TI)/Echo time (TE)=2300/900/2.98 ms, flip angle=9 degrees, voxel dimensions=1 mm isotropic, 32 slices). Functional images were acquired with a susceptibility weighted EPI sequence (TR/TE=2000/30 ms, flip angle=75 degrees, voxel dimensions=3×3×3.5 mm, 192 slices).

#### Data Analysis

Functional MRI data were preprocessed and then analyzed using the general linear model (GLM) for event-related designs in SPM8 (Welcome Department of Imaging Neuroscience, London, UK; http://www.fil.ion.ucl.ac.uk/spm). During preprocessing, all functional volumes were realigned to the mean image, co-registered to the structural T1 image, corrected for slice timing, normalized to the MNI EPI-template, and smoothed using an 8 mm FWHM Gaussian kernel. Statistical analyses were performed on a voxel wise basis across the whole-brain. At the first-level analysis, individual events were modelled by a standard synthetic hemodynamic response function (HRF) and six rigid-body realignment parameters were included as nuisance covariates when estimating statistical maps. Loss and Gain conditions were modeled as separate sessions within the same GLM. Two types of analysis were conducted to investigate the neural correlates of exploratory decision making.

### fMRI analysis 1: Exploration versus exploitation

The purpose of this model was to define regions-of-interest (ROIs) to be used in subsequent analyses. To elucidate brain regions engaged by decisions to explore vs. exploit, an event-related design was created with four event-types (exploratory decision, exploitation, response, and feedback) for each Gain/Loss condition separately. The first two event-types were time-locked to the presentation of the slot machines (decision onset), the response was locked to the button press, and the feedback was time-locked to the presentation of the outcome (feedback onset). Trials which could not be categorized as exploration or exploitation were included in a regressor-of-no-interest together with trials in which no response was provided.

### fMRI analysis 2: Parametric modulation of decisions by differences in expected value and outcome uncertainty, and by prediction error at feedback

To determine brain regions tracking trial-by-trial differences in expected value (*ΔQ*) and outcome uncertainty (*ΔT*), as well as prediction errors (*δ*), an event-related design was created with three event-types (decision onset, response onset, feedback onset) time-locked to the presentation of the slot machines, the button press, and the presentation of outcomes, respectively. In each trial, *ΔQ* was defined as the difference in expected value between the average expected value of the two rejected machines and the selected machine. Similarly, *ΔT* was defined as the difference in uncertainty between the average uncertainty for the two rejected machines and the selected machine. These two parameters were added as parametric modulators at decision onset, while *δ* was added as a parametric modulator of the feedback onset. Trials in which no response was provided were included in a regressor-of-no-interest.

### Regions of interest (ROIs)

#### Exploratory decision making

The dorsal anterior cingulate cortex (dACC), the anterior insulae (aINS), and the frontopolar cortex (FPC) are engaged by different aspects of uncertainty and by decisions to explore (Badre et al. 2012; Blanchard and Gershman 2018; Daw et al. 2006; Laureiro-Martinez et al. 2015; Grupe and Nitschke 2013), while exploitation activates the ventromedial prefrontal cortex (vmPFC; (Laureiro-Martinez et al. 2015)). Our fMRI analyses therefore focus on these brain regions. A dACC ROI was created using the WFU Pickatlas toolbox (Maldjian et al. 2003) in combination with a previously described procedure (Cascio, Konrath, and Falk 2015; Wang et al. 2016): First, the union of Broadmann areas 24 and 32, and AAL masks of the cingulate cortex (anterior, middle, posterior; dilated to 2mm) was created. Next, Broadmann areas 8 and 9 were subtracted from this union, and the resulting ROI was further restricted by a cuboid with bounds x = [−16 16], y = [0 33], z = [6 52]. The bilateral aINS was defined by the left and right insula ROIs of the WFU pickatlas toolbox. The FPC ROI was defined as the bilateral Area Fp1 of the Anatomy toolbox (Eickhoff et al. 2005; Bludau et al. 2014). The vmPFC ROI was defined as a 10 mm radius sphere centered on x=−3, y=42, z=−15, calculated from a recent meta-analysis investigating the role of the vmPFC/orbitofrontal cortex in representing a neural common currency for choice (Levy and Glimcher 2012). These ROIs were combined into one ROI mask.

#### Prediction error

The Nucleus Accumbens (NAcc) encodes prediction errors (Aberg, Doell, and Schwartz 2016a; Cohen 2007). The NAcc ROI was obtained from a recent study trying to delineate the shell and core subregions of the NAcc (Baliki et al. 2013).

### Statistical analysis

To investigate how the neural correlates of exploratory decision making are modulated by trait anxiety, the fMRI data was analyzed in two steps. This analysis follows recent recommendations for correlating inter-individual factors, such as personality traits, with fMRI data (Vul et al. 2009).

First, fMRI analysis 1 was used in combination with the dACC/aINS/FPC/vmPFC ROI mask to identify group-level peak-voxel activity for the contrast between exploratory decisions and exploitation (and vice versa). These activations were identified using a threshold of p=0.001 (uncorrected) and a minimum cluster size of five contiguous voxels. Small volume correction (SVC) using a threshold of p<0.05 Family-Wise Error Rate (FWER) for multiple comparisons were applied using the ROI mask. Importantly, because this analysis was performed across Gain/Loss condition and individuals, it allows an unbiased selection of brain coordinates for studying the differential contribution of Gain/Loss condition and modulations by trait anxiety.

Second, fMRI analysis 2 was used to extract beta parameter estimates (i.e. *β*_Δ*Q*_ and *β*_Δ*T*_) for the different parametric modulators (i.e. *ΔQ* and *ΔT*, respectively). Activity was extracted by averaging activity within 3 mm radius spheres centered on the coordinates of the peak voxels identified via fMRI analysis 1. The resulting data was then entered into repeated measures ANOVAs with one within-subject factor Condition (Gain, Loss) and continuous covariate Trait anxiety. Statistical thresholds were Bonferroni-corrected based on the number of ANOVAs conducted for each condition. To ensure that any significant result was not critically dependent on the 3mm radius of the sphere (i.e. by averaging the voxel intensity across five voxels), we repeated the same analyses using spheres with radiuses of 0mm (1 voxel), 5mm (19 voxels), and 10mm (137 voxels) respectively. The resulting p-values of these complementary analyses are presented together with the results of the main analyses that used 3mm radius spheres.

## Notes

### Competing Interest Statement

The authors have declared no competing interest.

### Summary of Updates

New analyses added, new data added.

