## Supplementary notes and figures for "A neural and behavioral tradeoff underlies exploratory decisions in normative anxiety"

### Supplementary Figures and Material

### Supplementary Note 1

We report the results of a behavioral prior study with 20 participants, independent and different from the participants in the main fMRI study. This study was conducted in order to test the hypothesis of whether anxiety affects exploration and to design an appropriate model to explain how anxiety affects exploration.

As in the main fMRI study, participants performed a three-armed bandit task, but only in a loss-condition. The experiment was identical to the fMRI experiment with the following exceptions: the magnitude and mean of each of the three sine-waves was always 30 and 50 points respectively. The period lengths of the three sine-waves in a session were 100, 100, or 25 trials, respectively, and the sine-waves were phase-shifted by 0, 0.5, and 0.75 period lengths.

Using the same analysis as described in the main text, we observed a positive correlation between the proportion of exploratory decisions and trait anxiety (r=0.485, p=0.030, two-tailed Pearson correlation; Supplementary Figure 1A). This was the main finding that led us to perform the fMRI experiment in which the result was independently replicated. Further, there was an inverse trend between overall performance and exploration (Supplementary Figure 1B), potentially due to the difference in magnitude between the fMRI (20) and pilot (30) study, making the task in the pilot study relatively easier and less exploration required. Based on previous research we designed different computational models (exactly the same models as described in the main article), fitted them to experimental data, and compared their fits using Bayesian Model Selection (BMS) with AICc as model evidence. As in the main article, BMS revealed that the model most likely to have generated the observed behavior was the model with one value learning rate ($\alpha_Q$) and decision weights for expected-value ($\beta_Q$), outcome-uncertainty ($\beta_T$), and random-switching ($\beta_S$; Exceedance probability=0.999; Supplementary Figure 1C). This is therefore an independent validation of the chosen model. To further validate this model, we used the fitted parameters to generate model-predicted exploration rates, and this result reproduced the positive correlation between trait anxiety and exploration observed in behavior (r=0.492, p=0.027, two-tailed Pearson correlation; Supplementary Figure 1D), again similar to and independently of the fMRI experiment. Finally, trait anxiety showed a positive correlation with the fitted model parameter related to uncertainty-reduction ($\beta_T$; r=0.536, p=0.015, two-tailed Pearson correlation; Supplementary Figure 1E), again validating the main fMRI experiment. There was a trend of inverse correlation between trait anxiety and the decision weight for expected-value ($\beta_Q$; r=-0.167, p=0.48, two-tailed Pearson correlation; Supplementary Figure 1F). We therefore hypothesized that a larger sample study would replicate the results and strengthen the trends, as indeed found in our main fMRI study reported in the manuscript.

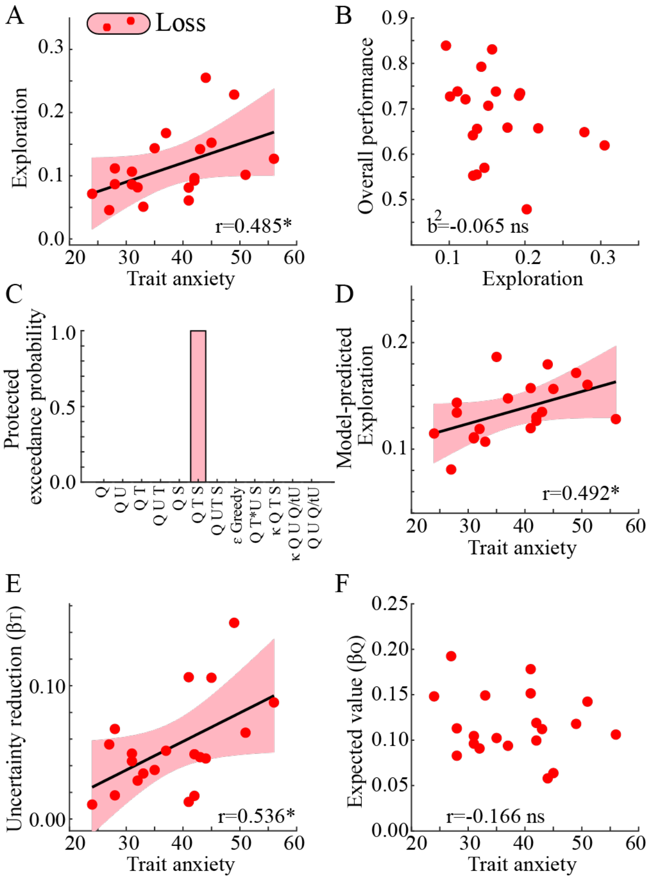

**Supplementary Figure 1.** Results from the preliminary (behavior-only) study. **A.** Like in the main fMRI experiment, exploration increased linearly with increased levels of trait anxiety. **B.** Overall performance, defined as the correlation between maximal and actual outcomes across all trials, seems to be inversely correlated with exploration. **C.** Protected exceedance probabilities for the different models. Similar to the main fMRI experiment, the QTS model was the most likely model here as well. **D.** The average model-predicted probability of exploration correlated positively with trait anxiety, therefore validating the selected model. **E.** Like in the main fMRI experiment, the decision weight estimating reduction of outcome uncertainty ($\beta_T$) was positively correlated with trait anxiety. **F.** The decision weight estimating the influence of expected value ($\beta_Q$) seems to be inversely correlated with trait anxiety. *p<0.05; ns p>0.05.

### Supplementary Note 2

To test for a significant difference between Loss and Gain conditions regarding the quadratic relationship between Exploration and Overall performance (i.e. $\Delta b^2_{Explore}$), a Monte-Carlo randomization procedure was applied. A null-distribution of differences was created by switching data points between Loss and Gain conditions within a subject (i.e. $\Delta b^2_{Explore, null}$). In each of 10'000 shuffles, the data from a random number of participants was switched between Loss and Gain conditions. For each of the shuffles, two nonlinear regressions were performed with intercept and quadratic term $b^2_{Explore}$, one for shuffled Loss ($b^2_{Explore, Loss}$) and one for shuffled Gain ($b^2_{Explore, Loss}$). The resulting difference in $b^2_{Explore}$ between Loss and Gain conditions were added to $\Delta b^2_{Explore, null}$. Finally, the proportion of samples whose absolute value was larger or equal to the observed $\Delta b^2_{Explore}$ was used to calculate a two-tailed p-value. The actually observed difference was -0.093, which did not differ significantly from the null-distribution of differences (null mean=-0.040, p=0.198). The null-distribution of differences is shown in Supplementary Figure 2.

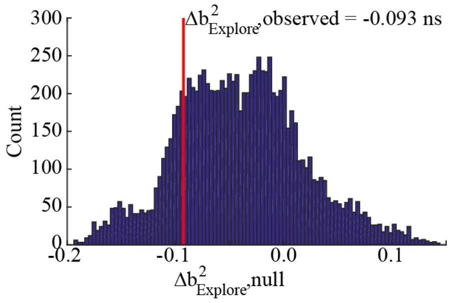

**Supplementary Figure 2.** Difference in the quadratic relationship between Exploration and Overall performance between Gain and Loss conditions. **A.** The observed difference in $\Delta b^2_{Explore}$ (red line) did not differ significantly from the null-distribution of differences $\Delta b^2_{Explore, null}$ (blue bars). ns p>0.05.

### Supplementary Note 3

To test for a significant difference between Loss and Gain conditions regarding the quadratic relationship between Trait anxiety and Overall performance, a Monte-Carlo randomization procedure was applied (see Supplementary Note 1). In brief, the actually observed difference was -0.483, which differed significantly from the null-distribution of differences (null mean=-0.258, p=0.032). The null-distribution of differences is shown in Supplementary Figure 3.

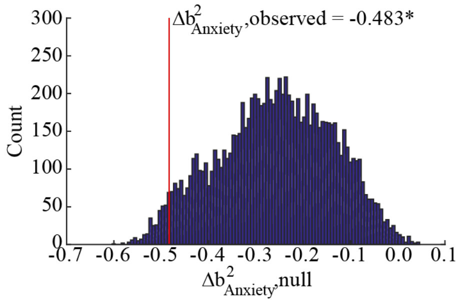

**Supplementary Figure 3.** Difference in the quadratic relationship between Trait anxiety and Overall performance between Gain and Loss conditions. **A.** The observed difference in $\Delta b^2_{Anxiety}$ (red line) differed significantly from the null-distribution of differences $\Delta b^2_{Anxiety,\ null}$ (blue bars). *p<0.05.

### Supplementary Note 4

A mediation analysis tested to what extent Exploration mediated the non-linear correlation between Trait anxiety and Overall performance (see Methods). The predictive value of Trait anxiety (on Overall performance) was significantly decreased ($\Delta b^2_{Anxiety} < 0$) across conditions, as well as separately in both the Loss and in the Gain condition, when Exploration was added to the regression analysis (Supplementary Table 1, Supplementary Figure 4). These results indicate that Exploration mediates the nonlinear relationship between anxiety and overall performance (Hayes & Rockwood, 2017).

***Supplementary Table 1. Mediation analysis.***

| | Predictor | β±STD | p-value |
|---|---|---|---|
| **Average across conditions** | | | |
| Trait anxiety only | $b^2_{Anxiety1}$ | -0.591 ± 0.136 | <0.001 |
| Exploration & Trait anxiety | $b^2_{Explore}$ | -0.081 ± 0.098 | 0.283 |
| | $b^2_{Anxiety2}$ | -0.440 ± 0.229 | 0.066 |
| | $\Delta b^2_{Anxiety} = b^2_{Anxiety1} - b^2_{Anxiety2}$ | -0.152 | <0.001* |
| | $\Delta b^2_{Anxiety, null}$ | -0.0006 ± 0.026 | |
| **Loss condition** | | | |
| Trait anxiety only | $b^2_{Anxiety1}$ | -0.712 ± 0.111 | <0.001 |
| Exploration & Trait anxiety | $b^2_{Explore}$ | -0.099 ± 0.090 | 0.283 |
| | $b^2_{Anxiety2}$ | -0.523 ± 0.205 | 0.017 * |
| | $\Delta b^2_{Anxiety} = b^2_{Anxiety1} - b^2_{Anxiety2}$ | -0.190 | <0.001* |
| | $\Delta b^2_{Anxiety, null}$ | -0.0006 ± 0.024 | |
| **Gain condition** | | | |
| Trait anxiety only | $b^2_{Anxiety1}$ | -0.228 ± 0.173 | 0.198 |
| Exploration & Trait anxiety | $b^2_{Explore}$ | -0.213 ± 0.136 | 0.129 |
| | $b^2_{Anxiety2}$ | 0.032 ± 0.236 | 0.893 |
| | $\Delta b^2_{Anxiety} = b^2_{Anxiety1} - b^2_{Anxiety2}$ | -0.260 | <0.001* |
| | $\Delta b^2_{Anxiety, null}$ | -0.0005 ± 0.030 | |

Trait anxiety only: only trait anxiety was used as a non-linear (quadratic) predictor of overall performance. Exploration & Trait anxiety: both trait anxiety and proportion of exploratory decisions were used a non-linear (quadratic) predictors of overall performance. $b^2_{Anxiety}$ denotes the standardized predictive value of Trait anxiety, while $b^2_{Explore}$ denotes the standardized predictive value of Exploration. $\Delta b^2_{Anxiety}$ denotes the difference in predictive value of Trait anxiety without and with Exploration, and its statistical significance was determined using a Monte-Carlo randomization procedure. $\Delta b^2_{Anxiety, null}$ denotes the null-distribution obtained from the Monte-Carlo randomization procedure. STD is the standard deviation of the mean. * Statistically significant effect of mediation, i.e. p-value<0.05.

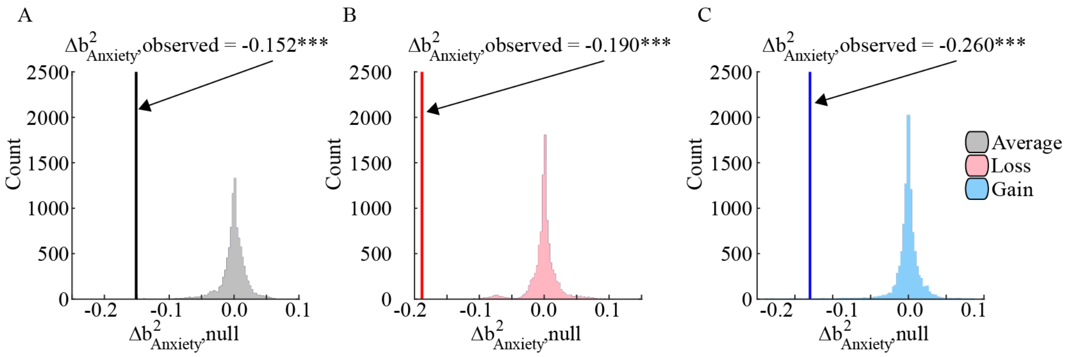

**Supplementary Figure 4.** Change in the quadratic relationship between Trait anxiety and Overall performance when controlling for Exploration. The observed difference in $\Delta b^2_{Anxiety}$ (indicated by vertical lines) differed significantly from the null-distribution of differences $\Delta b^2_{Anxiety,\ null}$ across conditions (**A**), and separately in the Loss condition **(B)** and in the Gain condition **(C)**. *p<0.05.

### Supplementary Note 5

The QTS model provided the most parsimonious fit to behavior, as determined by comparing the goodness of fit using the AICc in a Bayesian Model Selection procedure (BMS; see Fig.3A of the main text). Because the Bayes Information Criterion (BIC) provides a larger penalty for complex models, we also compared model fits using the BIC as model evidence and repeated the BMS procedure. Collapsed across Loss and Gain conditions, the QTS model again provided the most parsimonious fit to behavior (Protected exceedance probability=0.947; Supplementary Figure 5A). Performing the same analysis separately for Loss and Gain conditions revealed that the QTS model was most likely to have generated behavior in both conditions (Protected exceedance probability in Loss=0.747: Supplementary Figure 5B; Gain=0.492: Supplementary Figure 5C). Because the current implementation of BMS only provides approximate results when comparing more than two models, and because the exceedance probabilities were similar for the Q and QTS models in the Gain condition, we repeated the BMS procedure using only the BIC of the Q and QTS models as input. The result showed a higher likelihood for the QTS model in both Gain and Loss conditions (Protected exceedance probability in Loss=0.937; Gain=0.801; see insets in Supplementary Figure 5 B,C).

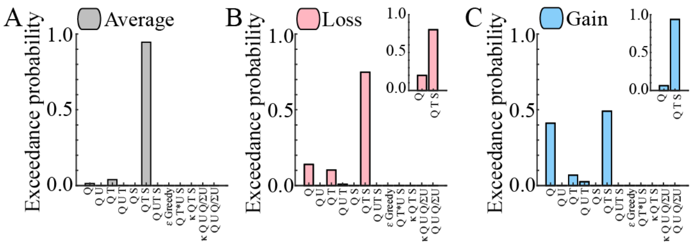

**Supplementary Figure 5.** Protected exceedance probabilities obtained using the Bayesian Information Criterion (BIC) as model evidence in the Bayesian Model Selection procedure. The different models collapsed across Loss and Gain conditions (**A**), and separately in the Loss (**B**) and in the Gain (**C**) condition. The insets in **A** and **B** show the BMS results when comparing only the Q and the QTS models.

Fitted parameters for each of the tested models are displayed separately for Gain and Loss conditions in Supplementary Table 2.

**Supplementary Table 2. Model fits.**

| Model | k | -LLE | AICc | $\alpha_Q$ | $\beta_Q$ | $\alpha_U$ | $\beta_U$ | $\beta_T$ | $\beta_S$ | $\varepsilon$ | $\beta_{tU}$ | $\beta_{U*T}$ | $Q_{ini}$ | $\sigma_0$ | $\sigma_{ini}$ |
|---|---|---|---|---|---|---|---|---|---|---|---|---|---|---|---|
| **Loss condition** | | | | | | | | | | | | | | | |
| Q | 2 | 102.7 | 209.49 | 0.95 | 0.12 | - | - | - | - | - | - | - | - | - | - |
| | | ± 7.15 | ± 14.30 | ± 0.02 | ± 0.01 | | | | | | | | | | |
| Q U | 4 | 104.78 | 217.78 | 0.95 | 0.12 | 0.33 | -0.17 | - | - | - | - | - | - | - | - |
| | | ± 6.97 | ± 13.94 | ± 0.02 | ± 0.01 | ± 0.07 | ± 0.09 | | | | | | | | |
| Q T | 3 | 99.08 | 204.30 | 0.94 | 0.14 | - | - | 0.04 | - | - | - | - | - | - | - |
| | | ± 7.19 | ± 14.39 | ± 0.02 | ± 0.01 | | | ± 0.01 | | | | | | | |
| Q S | 3 | 98.74 | 203.60 | 0.95 | 0.11 | - | - | - | -0.38 | - | - | - | - | - | - |
| | | ± 6.75 | ± 13.51 | ± 0.02 | ± 0.01 | | | | ± 0.10 | | | | | | |
| Q U T | 5 | 101.11 | 212.55 | 0.93 | 0.14 | 0.37 | -0.13 | 0.05 | - | - | - | - | - | - | - |
| | | ± 6.96 | ± 13.92 | ± 0.02 | ± 0.01 | ± 0.08 | ± 0.09 | ± 0.01 | | | | | | | |
| **Q T S** | **4** | **93.12** | **194.46** | **0.92** | **0.09** | **-** | **-** | **0.13** | **-0.76** | **-** | **-** | **-** | **-** | **-** | **-** |
| | | **± 6.46** | **± 12.92** | **± 0.02** | **± 0.02** | | | **± 0.01** | **± 0.13** | | | | | | |
| Q U T S | 6 | 95.96 | 204.37 | 0.91 | 0.14 | 0.34 | -0.13 | 0.10 | -0.66 | - | - | - | - | - | - |
| | | ± 6.17 | ± 12.34 | ± 0.02 | ± 0.01 | ± 0.08 | ± 0.09 | ± 0.02 | ± 0.13 | | | | | | |
| ε- | 2 | 112.65 | 229.37 | 0.87 | - | - | - | - | - | 0.21 | - | - | - | - | - |
| Greedy | | ± 6.62 | ± 13.23 | ± 0.02 | | | | | | ± 0.03 | | | | | |
| Q T*U S | 5 | 95.42 | 200.84 | 0.92 | 0.14 | 0.39 | 0.02 | - | - | - | - | -0.76 | - | - | - |
| | | ±5.93 | ±5.93 | ±0.02 | ±0.01 | ±0.06 | ±0.01 | | | | | ±0.16 | | | |
| κ-Q T S | 6 | 139.24 | 290.47 | - | 0.16 | - | - | -0.01 | -1.53 | - | - | - | -7.23 | 4 | 3.87± |
| | | ±6.09 | ±12.19 | | ±0.03 | | | ±0.02 | ±0.11 | | | | ±3.47 | | 0.54 |
| κ-Q U tU | 6 | 184.83 | 381.66 | - | 0.44 | - | 1.05 | - | - | - | -0.36 | - | -38.46 | 4 | 5.38 |
| | | ±57.63 | ±15.27 | | ±0.10 | | ±0.57 | | | | ±0.10 | | ±6.71 | | ±0.66 |
| Q T S tU | 5 | 94.99 | 201.98 | 0.89 | 0.06 | 0.35 | - | 0.11 | -0.71 | - | 2.01 | - | - | - | - |
| | | ±6.22 | ±12.44 | ±0.03 | ±0.03 | ±0.06 | | ±0.02 | ±0.14 | | ±0.57 | | | | |
| **Gain condition** | | | | | | | | | | | | | | | |
| Q | 2 | 93.66 | 191.38 | 0.91 | 0.14 | - | - | - | - | - | - | - | - | - | - |
| | | ± 7.51 | ± 15.01 | ± 0.03 | ± 0.01 | | | | | | | | | | |
| Q U | 4 | 95.69 | 199.59 | 0.85 | 0.16 | 0.30 | -0.09 | - | - | - | - | - | - | - | - |
| | | ± 7.44 | ± 14.88 | ± 0.04 | ± 0.02 | ± 0.06 | ± 0.08 | | | | | | | | |
| Q T | 3 | 89.27 | 184.68 | 0.89 | 0.16 | - | - | 0.04 | - | - | - | - | - | - | - |
| | | ± 7.21 | ± 14.43 | ± 0.03 | ± 0.01 | | | ± 0.02 | | | | | | | |
| Q S | 3 | 87.92 | 181.97 | 0.90 | 0.13 | - | - | - | -0.47 | - | - | - | - | - | - |
| | | ± 6.87 | ± 13.75 | ± 0.03 | ± 0.01 | | | | ± 0.12 | | | | | | |
| Q U T | 5 | 91.35 | 193.02 | 0.87 | 0.18 | 0.24 | -0.04 | 0.05 | - | - | - | - | - | - | - |
| | | ± 7.04 | ± 14.08 | ± 0.03 | ± 0.02 | ± 0.06 | ± 0.10 | ± 0.02 | | | | | | | |
| **Q T S** | **4** | **82.61** | **173.44** | **0.85** | **0.16** | **-** | **-** | **0.10** | **-0.95** | **-** | **-** | **-** | **-** | **-** | **-** |
| | | **± 6.52** | **± 13.05** | **± 0.04** | **± 0.01** | | | **± 0.02** | **± 0.14** | | | | | | |
| Q U T S | 6 | 85.25 | 182.97 | 0.82 | 0.17 | 0.25 | -0.10 | 0.11 | -0.89 | - | - | - | - | - | - |
| | | ± 6.37 | ± 12.74 | ± 0.04 | ± 0.02 | ± 0.06 | ± 0.10 | ± 0.03 | ±0.14 | | | | | | |
| ε- | 2 | 102.46 | 208.98 | 0.83 | - | - | - | - | - | 0.20 | - | - | - | - | - |
| Greedy | | ± 7.38 | ±14.75 | ±0.04 | | | | | | ± 0.03 | | | | | |
| Q T*U S | 4 | 85.99 | 181.97 | 0.82 | 0.17 | 0.37 | 0.02 | - | - | - | -1.00 | - | - | - | - |
| | | ±6.52 | ±11.86 | ±0.04 | ±0.02 | ±0.06 | ±0.01 | | | | ±0.15 | | | | |
| κ-Q T S | 6 | 129.68 | 271.37 | - | 0.17 | - | - | -0.00 | -1.76 | - | - | - | 73.69± | 4 | 5.22± |
| | | ±6.78 | ±13.56 | | ±0.04 | | | ±0.02 | ±0.12 | | | | 6.61 | | 0.71 |
| κ-Q U tU | 6 | 188.17 | 388.34 | - | 0.44 | - | 1.15 | - | - | - | -0.39 | - | 35.31 | 4 | 6.66 |
| | | ±4.28 | ±8.55 | | ±0.10 | | ±0.53 | | | | ±0.10 | | ±7.06 | | ±0.67 |
| Q T S tU | 4 | 85.47 | 182.95 | 0.82 | 0.07 | 0.38 | - | 0.11 | -0.96 | - | 1.78 | - | - | - | - |
| | | ±6.54 | ±13.08 | ±0.04 | ±0.03 | ±0.07 | | ±0.02 | ±0.13 | | ±0.57 | | | | |

k is the number of fitted parameters. -LLE is the negative log-likelihood. AICc is the Akaike Information Criterion corrected for small sample size. $\alpha_Q$ is the learning rate for the expecte-value. $\beta_Q$ is the decision weight for the expected-value. $\alpha_U$ is the learning rate for the outcome-variability. $\beta_U$ is the decision weight for the outcome-variability. $\beta_T$ is the decision weight for the outcome-uncertainty. $\beta_S$ is the decision weight for random switching. ε is the exploration probability for the ε-Greedy model. $\beta_{tU}$ is the decision weight for expected-value/total uncertainty-variability. $\beta_{U*T}$ is the decision weight for the outcome-uncertainty scaled by the outcome-variability. $Q_{ini}$ is the initial expected-value. $\sigma_{ini}$ is the initial outcome-variance. $\sigma_0$ was held constant at 4 for model degeneracy. The most parsimonious model in each condition is highlighted with bold font. Mean ± SEM.

### Supplementary Note 6

To determine the modulation of decision weights by Gain/Loss condition and trait anxiety, separate repeated measures ANOVAs with factor Condition (Gain, Loss) and continuous covariate Trait anxiety were conducted for each of the individually fitted decision weights for the QTS model. The significance threshold was Bonferroni-corrected to 0.0125 (0.05/4) for each of the four ANOVAs, and the results are reported in Supplementary Table 3 and displayed in Fig.4A-D and Supplementary Figure 6A-D.

**Supplementary Table 3.** Repeated measures ANOVAs for the fitted parameters $\beta_Q$, $\beta_T$, $\beta_S$, and $\alpha_Q$ as a function of Gain/Loss condition and Trait anxiety.

| | Sum of Squares | df | Mean Square | F | p-value (uncorrected) | $\eta_p^2$ |
|---|---|---|---|---|---|---|
| $\boldsymbol{\beta_Q}$ | | | | | | |
| Intercept | 1.174 | 1 | 1.174 | 240.26 | <.001 * | |
| Trait anxiety | 0.041 | 1 | 0.041 | 8.407 | 0.008 * | 0.244 |
| Error | 0.127 | 26 | 0.005 | | | |
| Condition | 0.007 | 1 | 0.007 | 3.015 | 0.094 | 0.104 |
| Trait anxiety x Condition | 0.0006 | 1 | 0.0006 | 0.250 | 0.622 | 0.010 |
| Error | 0.064 | 26 | 0.003 | | | |
| $\boldsymbol{\beta_T}$ | | | | | | |
| Intercept | 0.483 | 1 | 0.483 | 63.424 | <.001 * | |
| Trait anxiety | 0.079 | 1 | 0.079 | 10.358 | 0.003 * | 0.285 |
| Error | 0.198 | 26 | 0.008 | | | |
| Condition | <0.001 | 1 | <0.001 | <0.001 | 0.994 | <0.001 |
| Trait anxiety x Condition | <0.001 | 1 | <0.001 | 0.028 | 0.869 | 0.001 |
| Error | 0.134 | 26 | 0.005 | | | |
| $\boldsymbol{\beta_S}$ | | | | | | |
| Intercept | 40.638 | 1 | 40.638 | 43.205 | <.001 * | |
| Trait anxiety | 0.150 | 1 | 0.150 | 0.160 | 0.693 | 0.006 |
| Error | 24.455 | 26 | 0.941 | | | |
| Condition | 0.502 | 1 | 0.502 | 4.082 | 0.054 | 0.136 |
| Trait anxiety x Condition | 0.259 | 1 | 0.259 | 2.107 | 0.159 | 0.075 |
| Error | 3.195 | 26 | 0.123 | | | |
| $\boldsymbol{\alpha_Q}$ | | | | | | |
| Intercept | 43.814 | 1 | 43.814 | 1923.1 | <.001 * | |
| Trait anxiety | 0.059 | 1 | 0.059 | 2.586 | 0.120 | 0.091 |
| Error | 0.592 | 26 | 0.023 | | | |
| Condition | 0.052 | 1 | 0.052 | 2.817 | 0.105 | 0.098 |
| Trait anxiety x Condition | 0.073 | 1 | 0.073 | 3.947 | 0.058 | 0.132 |
| Error | 0.480 | 26 | 0.019 | | | |

$\beta_Q$ is the decision weight for the expected outcome. $\beta_T$ is the decision weight for the outcome uncertainty. $\beta_S$ is the decision weight for the random switching. $\alpha_Q$ is the learning rate for the expected outcome. df:

Degrees of Freedom. F: F-statistic.

$\eta_p^2$ : Partial eta-squared. * indicates significant p-value at the Bonferroni-corrected threshold (α=0.0125).

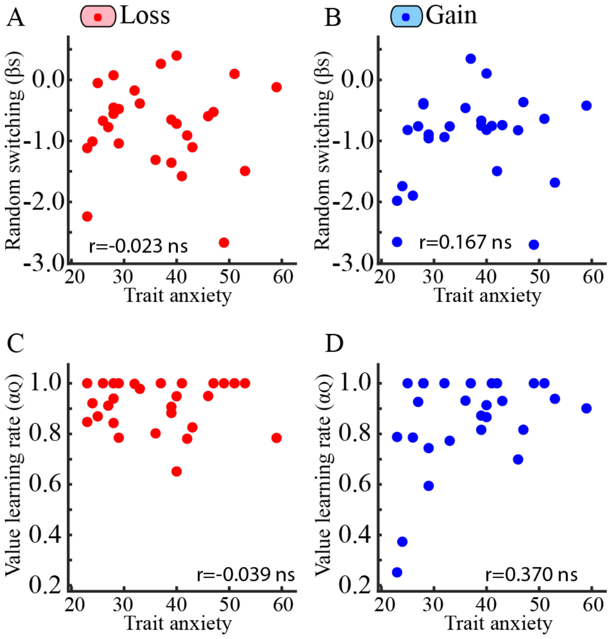

**Supplementary Figure 6.** Individually fitted model parameters versus Trait anxiety and Gain/Loss condition. **A-B.** Decision weights estimating the influence of randomly switching machines ($\beta_S$) did not correlate with trait anxiety. **C-D.** The value learning rate ($\alpha_Q$) did not correlate with trait anxiety. ns p>0.05.

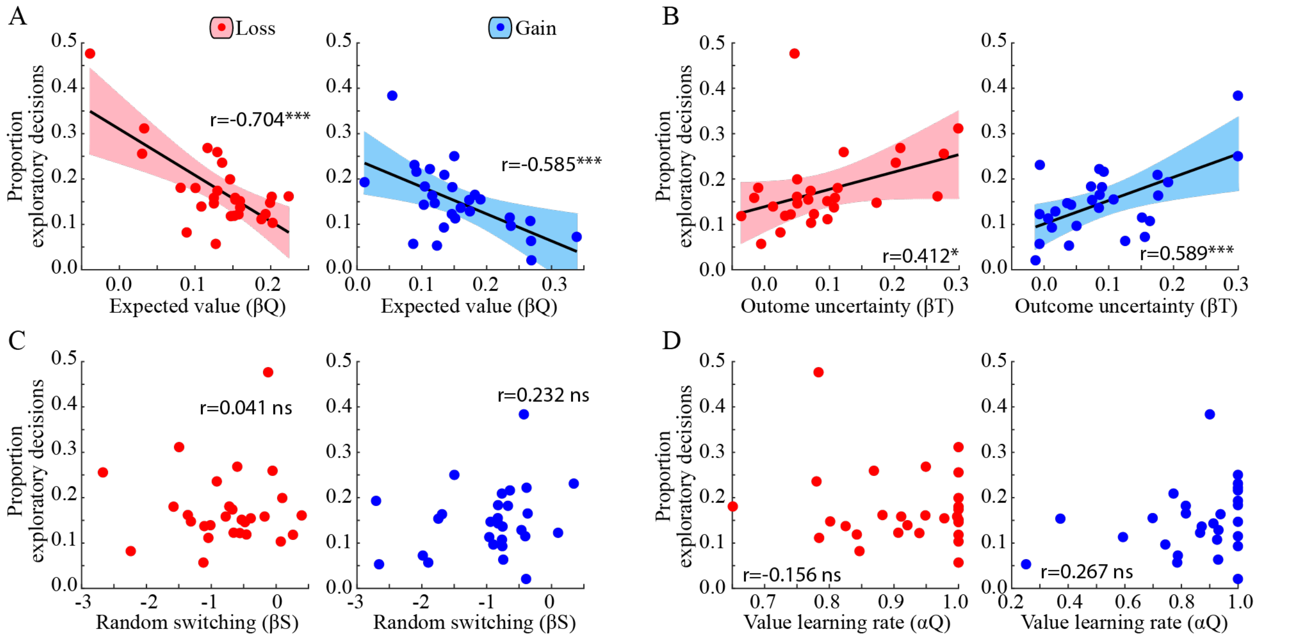

**Supplementary Figure 7.** Individually fitted model parameters versus the proportion of exploratory decisions. **A.** The proportion of exploratory decisions correlated negatively with the decision weight estimating the influence from expected value (i.e. $\beta_Q$) in both the Loss and the Gain condition. **B.** The proportion of exploratory decisions correlated positively with the decision weight estimating uncertainty reduction (i.e. $\beta_T$) in both conditions. **C.** Exploration was not modulated by the inter-individual tendency to randomly switch machines (i.e. $\beta_S$). **D.** Exploration was not related to the value learning rate (i.e. $\alpha_Q$).***p<0.001; *p<0.05; ns p>0.05.

### Supplementary Note 7

To provide a more robust assessment of the decision variables underpinning increased exploration in trait anxiety, we compared the proportion of exploratory decisions and decision weights derived from a Kalman-filter model using a dynamic learning rate, but with the same decision weights as the most parsimonious QTS model with a constant learning rate (see Methods).

Using the same model-based definition of Exploration, i.e. the proportion of switches to non-optimal machines, and the same ANOVA as in the main text (Trait anxiety as continuous covariate, and Gain/Loss condition as within-subject factor), revealed a significant and positive correlation between Trait anxiety and Exploration (F(1, 26)=29.336, p<0.001; Supplementary Fig.8A), while no other effects or interactions were significant (all p-values>0.11). Testing the model-fitted decision weights related to uncertainty-reduction ($\beta_{\mathrm{T}}$) using the ANOVA with the same factors as above, revealed a positive and significant main effect of Trait anxiety (F(1, 26)=6.877, p=0.014; Supplementary Figure 8B), while no other effects or interactions were significant (all p-values>0.608). By contrast, the same ANOVA for the expected-value factor ($\beta_{\mathrm{T}}$) revealed no significant effects (all p-values > 0.06; Supplementary Figure 8C).

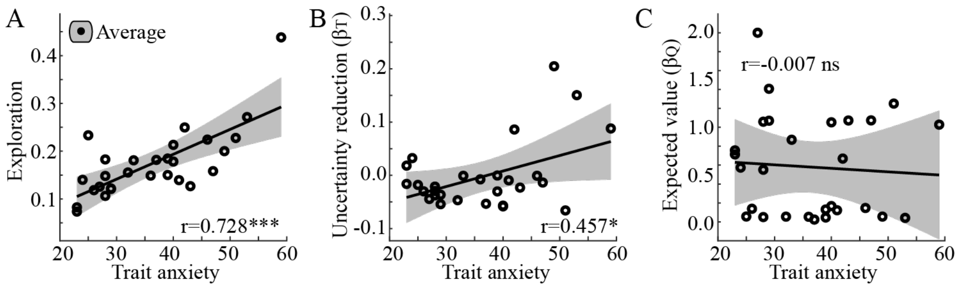

**Supplementary Figure 8. A.** Exploration, the proportion of exploratory decisions, increased linearly with increased levels of trait anxiety across conditions. **B.** Decision weights estimating reduction of outcome uncertainty ($\beta_T$) was positively correlated with trait anxiety across conditions. **C.** Decision weights estimating the influence of expected value ($\beta_Q$) did not correlate significantly with trait anxiety. p<0.001; *p<0.05; ns p>0.05.

### Supplementary Note 8

The prediction error signal δ collapsed across participants and Gain/Loss conditions correlated significantly with activity in the bilateral NAcc [Supplementary Figure 8; Right NAcc: peak voxel MNI xyz = 9 11 -5, p<0.001 (FWE, SVC); Left NAcc: peak voxel MNI xyz = -9 11 -5, p<0.001 (FWE, SVC)]. Beta parameter estimates of δ (i.e. $\beta_\delta$), extracted using 3mm radius spheres centered on these coordinates, were entered into two repeated measures ANOVAs with factor Condition (Gain, Loss) and Trait anxiety as continuous covariate. For the two ANOVAs conducted, the statistical threshold was Bonferroni-corrected to 0.025 (0.05/2). There were no significant effects of Condition, Trait anxiety, nor their interaction (Supplementary Figure 9; Supplementary Table 4).

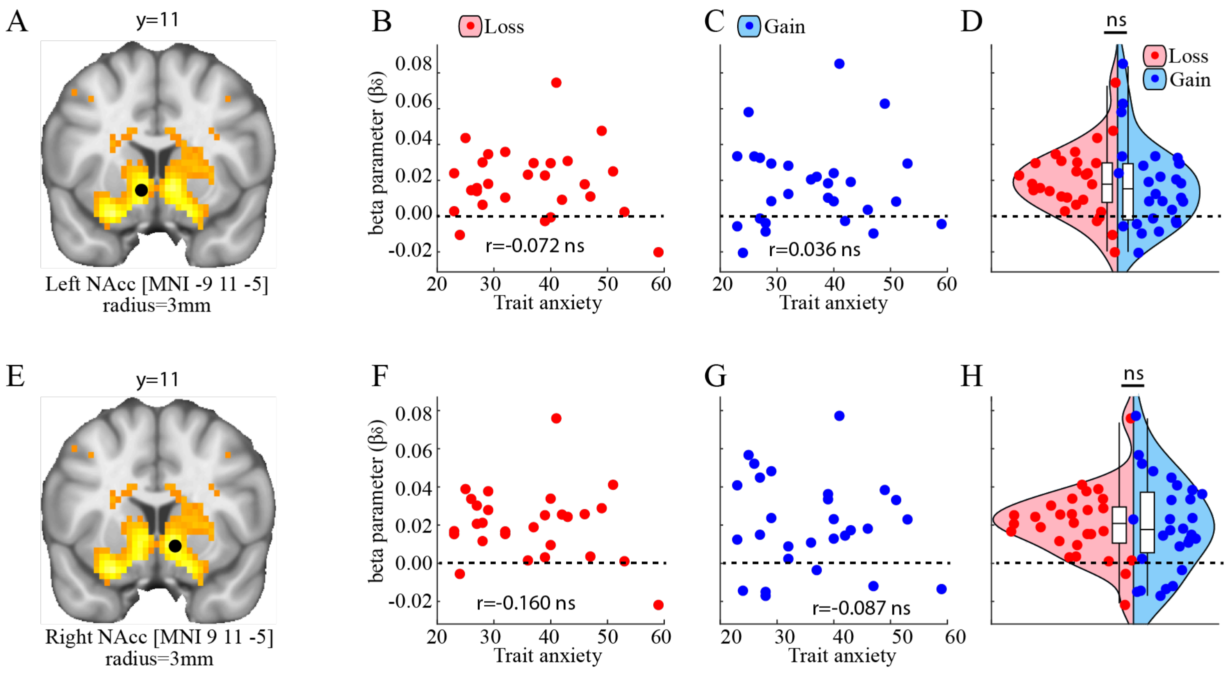

**Supplementary Figure 9.** The neuronal representation (i.e. $\beta_\delta$) of the prediction error δ. BOLD signal in the Nucleus Accumbens (NAcc) was significantly modulated by δ in both the left (**A-D**) and the right hemisphere (**E-H**), but $\beta_\delta$ was not modulated by trait anxiety nor Gain/Loss condition. ns $p > 0.05$. For visualization purposes, the brain activation is displayed using an uncorrected threshold of p=0.001. Beta parameters were extracted from 3mm radius spheres (indicated by black spheres) centered on the coordinates reported in the left-most column.

**Supplementary Table 4.** Repeated measures ANOVA for $\beta_\delta$ as a function of Gain/Loss conditions and Trait anxiety in the left and right NAcc.

| | Sum of Squares | df | Mean Square | F | p-value (uncorrected) |
|---|---|---|---|---|---|
| Left NAcc | | | | | |
| Intercept | 0.019 | 1 | 0.019 | 22.02 | <0.001 * |
| Trait anxiety | <0.0001 | 1 | <0.0001 | 0.005 | 0.946 |
| Error | 0.022 | 26 | 0.0009 | | |
| Condition | <0.0001 | 1 | <0.0001 | 0.39 | 0.537 |
| Trait anxiety x Condition | <0.0001 | 1 | <0.0001 | 0.62 | 0.437 |
| Error | 0.003 | 26 | 0.0001 | 0.44 | |
| Right NAcc | | | | | |
| Intercept | 0.023 | 1 | 0.023 | 29.25 | <0.001 * |
| Trait anxiety | 0.0003 | 1 | 0.0003 | 0.422 | 0.521 |
| Error | 0.021 | 26 | 0.0008 | | |
| Condition | <0.0001 | 1 | <0.0001 | 0.010 | 0.919 |
| Trait anxiety x Condition | <0.0001 | 1 | <0.0001 | 0.064 | 0.803 |
| Error | 0.0035 | 26 | 0.0001 | | |

NAcc = nucleus accumbens. * indicates that the p-value is significant at the Bonferroni-corrected threshold (α=0.025).

### Supplementary Note 9

To identify how BOLD signal is modulated by differences in expected values $\Delta Q$ between the average of the two rejected machines and the selected machine, beta parameter estimates of $\Delta Q$ (i.e. $\beta_{\Delta Q}$) were extracted using 3 mm radius spheres surrounding the peak voxels independently identified via the functional localizer that contrasted decisions to explore and exploit (see main text). Separate repeated measures ANOVAs were conducted for each of these ROIs, with within-subject factor Condition (Gain, Loss) and Trait anxiety as continuous covariate.

Five different ANOVAs were conducted (one for each of the five ROIs), thus the Bonferroni-corrected significance threshold was set to 0.01 (0.05/5). Extracted beta parameters for the different ROIs are shown in Supplementary Figure 9 and the results of the ANOVAs are presented in Supplementary Table 5. In brief, $\beta_{\Delta Q}$ was significantly and positively correlated with Trait anxiety in the dACC but not in the other ROIs (for an elaborate discussion of this results, see main text), nor did any of these ROIs show differential $\beta_{\Delta Q}$ between Gain/Loss conditions.

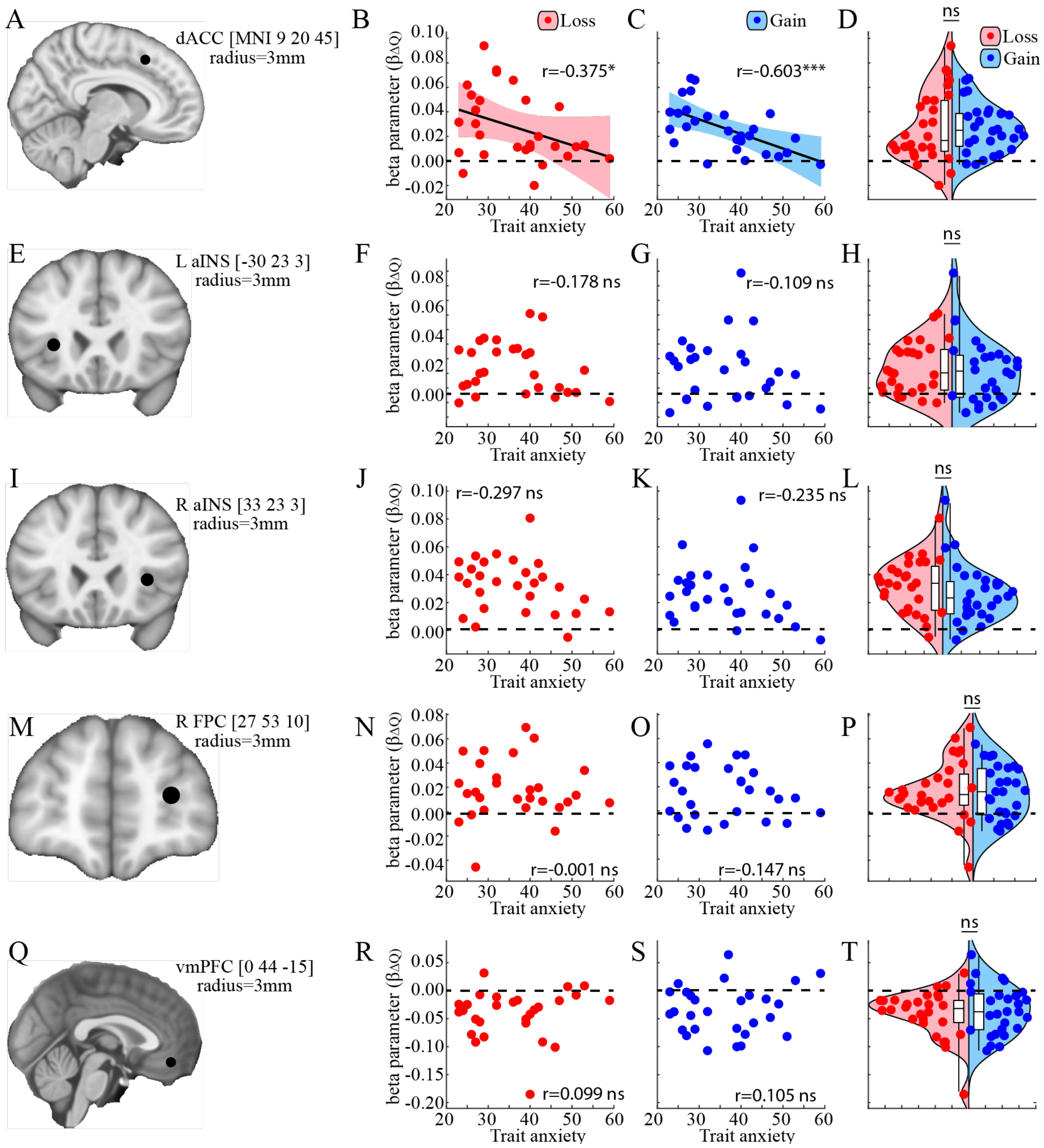

**Supplementary Figure 10.** Beta parameter estimates (i.e. $\beta_{\Delta Q}$) of expected value differences (i.e. $\Delta Q$) as a function of Gain/Loss condition and Trait anxiety. Beta parameters were extracted from 3mm radius spheres (indicated by black dots) centered on the coordinates reported in the left-most column. **A-C.** $\beta_{\Delta Q}$ correlated negatively with trait anxiety in the dACC in the Loss (**B**) and in the Gain condition (**C**), but did not differ between Gain/Loss conditions (**D**). **E-T.** $\beta_{\Delta Q}$ did not correlate with trait anxiety and did not differ between Gain/Loss conditions in the left aINS (**E-H**), the right aINS (**I-L**), the right FPC (**M-P**), or the vmPFC (**Q-T**). ***p<0.001; *p<0.05; ns p>0.05.

**Supplementary Table 5.** Repeated measures ANOVA for $\beta_{\Delta Q}$ as a function of Gain/Loss conditions and Trait anxiety in each of the a priori ROIs.

| | Main analysis | | | | | Robustness analysis p-value (uncorrected) | | |
|---|---|---|---|---|---|---|---|---|
| | Sum of Squares | df | Mean Square | F | p-value (uncorrected) r=3 | r=0 | r=5 | r=10 |
| vmPFC | | | | | | | | |
| Intercept | 0.081 | 1 | 0.081 | 29.933 | <.001 * | <.001 * | <.001 * | <.001 * |

| | | | | | | | | |
|---|---|---|---|---|---|---|---|---|
| Trait anxiety | 0.001 | 1 | 0.001 | 0.390 | 0.538 | 0.5138 | 0.5577 | 0.577 |
| Error | 0.070 | 26 | 0.003 | | | | | |
| Condition | 0.001 | 1 | 0.001 | 0.539 | 0.470 | 0.437 | 0.557 | 0.737 |
| Trait anxiety x Condition | <.001 | 1 | <.001 | 0.001 | 0.973 | 0.985 | 0.933 | 0.824 |
| Error | 0.030 | 26 | 0.001 | | | | | |
| <u>dACC</u> | | | | | | | | |
| Intercept | 0.041 | 1 | 0.041 | 60.594 | <.001 * | <.001 * | <.001 * | <.001 * |
| Trait anxiety | 0.007 | 1 | 0.007 | 10.242 | 0.004 * | 0.005 * | 0.004 * | 0.008 * |
| Error | 0.018 | 26 | <.001 | | | | | |
| Condition | <.001 | 1 | <.001 | 0.059 | 0.811 | 0.793 | 0.767 | 0.649 |
| Trait anxiety x Condition | <.001 | 1 | <.001 | 0.059 | 0.810 | 0.978 | 0.696 | 0.544 |
| Error | 0.008 | 26 | <.001 | | | | | |
| <u>Right FPC</u> | | | | | | | | |
| Intercept | 0.016 | 1 | 0.016 | 24.049 | <.001 * | <.001 * | <.001 * | <.001 * |
| Trait anxiety | <.001 | 1 | <.001 | 0.206 | 0.654 | 0.638 | 0.620 | 0.503 |
| Error | 0.017 | 26 | <.001 | | | | | |
| Condition | <.001 | 1 | <.001 | 0.125 | 0.726 | 0.749 | 0.680 | 0.626 |
| Trait anxiety x Condition | <.001 | 1 | <.001 | 0.319 | 0.577 | 0.583 | 0.565 | 0.538 |
| Error | 0.011 | 26 | <.001 | | | | | |
| <u>Left aINS</u> | | | | | | | | |
| Intercept | 0.029 | 1 | 0.029 | 46.286 | <.001 * | <.001 * | <.001 * | <.001 * |
| Trait anxiety | <.001 | 1 | <.001 | 0.642 | 0.430 | 0.481 | 0.427 | 0.396 |
| Error | 0.020 | 26 | <.001 | | | | | |
| Condition | <.001 | 1 | <.001 | 0.209 | 0.652 | 0.524 | 0.633 | 0.499 |
| Trait anxiety x Condition | <.001 | 1 | <.001 | 0.042 | 0.839 | 0.864 | 0.957 | 0.818 |
| Error | 0.005 | 26 | <.001 | | | | | |
| <u>Right aINS</u> | | | | | | | | |
| Intercept | 0.048 | 1 | 0.048 | 76.473 | <.001 * | <.001 * | <.001 * | <.001 * |
| Trait anxiety | <.001 | 1 | <.001 | 2.429 | 0.131 | 0.161 | 0.138 | 0.056 |
| Error | 0.020 | 26 | <.001 | | | | | |
| Condition | <.001 | 1 | <.001 | 3.040 | 0.093 | 0.091 | 0.095 | 0.077 |
| Trait anxiety x Condition | <.001 | 1 | <.001 | 0.039 | 0.844 | 0.976 | 0.876 | 0.683 |
| Error | 0.004 | 26 | <.001 | | | | | |

vmPFC = ventromedial prefrontal cortex. dACC=dorsal anterior cingulate cortex. FPC = right frontopolar cortex. aINS = anterior insula. r=0, r=3, r=5, r=10 refers to the radius of the sphere used to extract the beta parameters and corresponds to 1, 5, 19, and 137 voxels respectively. * indicates that the p-value is significant at the Bonferroni-corrected threshold (α=0.01).

### Supplementary Note 10

To test whether more anxious individuals displayed overall increased dACC activation during decision making, i.e. independent of any modulating factor, beta parameters ($\beta_{\text{Decision}}$) were extracted from 3mm spheres centered around the MNI coordinates xyz=9 20 45. A repeated measures ANOVA with factor Condition (Gain, Loss) and continuous covariate Trait anxiety revealed no effect or interaction with Trait anxiety (Supplementary Figure 10; Supplementary Table 6).

**Supplementary Table 6.** Repeated measures ANOVA for $\beta_{Decision}$ as a function of Gain/Loss conditions and Trait anxiety in each of the a priori ROIs.

| | Sum of Squares | df | Mean Square | F | p-value (uncorrected) r=3 |
|---|---|---|---|---|---|
| dACC | | | | | |
| Intercept | 39.884 | 1 | 39.884 | 50.916 | <0.001 |
| Trait anxiety | 0.021 | 1 | 0.021 | 0.027 | 0.870 |
| Error | 20.367 | 26 | 0.783 | | |
| Condition | 1.139 | 1 | 1.139 | 9.925 | 0.004 |
| Trait anxiety x Condition | 0.116 | 1 | 0.116 | 1.009 | 0.324 |
| Error | 2.984 | 26 | 0.115 | | |

dACC=dorsal anterior cingulate cortex.

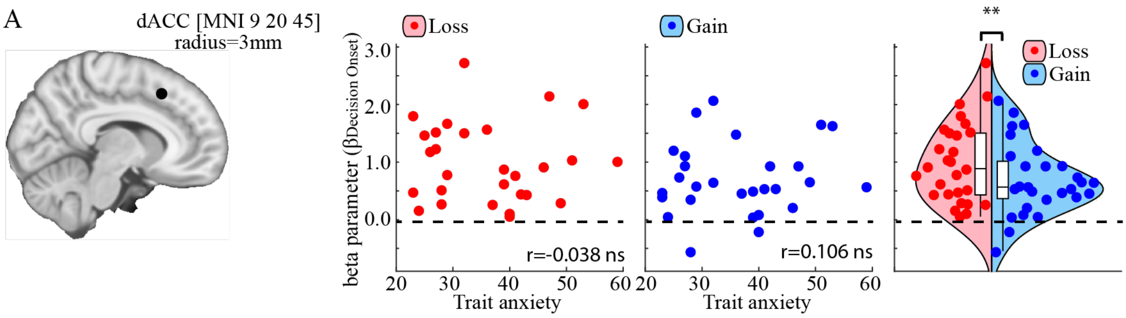

**Supplementary Figure 11.** BOLD signal at decision onset. **A-B.** Trait anxiety did not correlate with the BOLD response at decision onset in the Loss (**A**) or the Gain condition (**B**). ns p>0.05.

### Supplementary Note 11

To identify how BOLD signal is modulated by differences in expected values $\Delta T$ between the two rejected machines and the selected machine. Beta parameter estimates of $\Delta T$ (i.e. $\beta_{\Delta T}$) were extracted using 3 mm radius spheres surrounding the peak voxels independently identified via the differential activation between decisions to explore and exploit (see main text). Separate repeated measures ANOVAs were conducted for each of these ROIs, with within-subject factor Condition (Gain, Loss) and Trait anxiety as continuous covariate.

As in the previous paragraph, a Bonferroni-corrected significance threshold of 0.01 was adopted for each of five conducted ANOVAs. Extracted beta parameters are shown in Supplementary Figure 11 and the results of the ANOVAs are reported in Supplementary Table 7. In brief, there was a significant and negative correlation between $\beta_{\Delta T}$ and Trait anxiety bilaterally in the aINS but not in any other ROIs (see main text for an elaborate discussion).

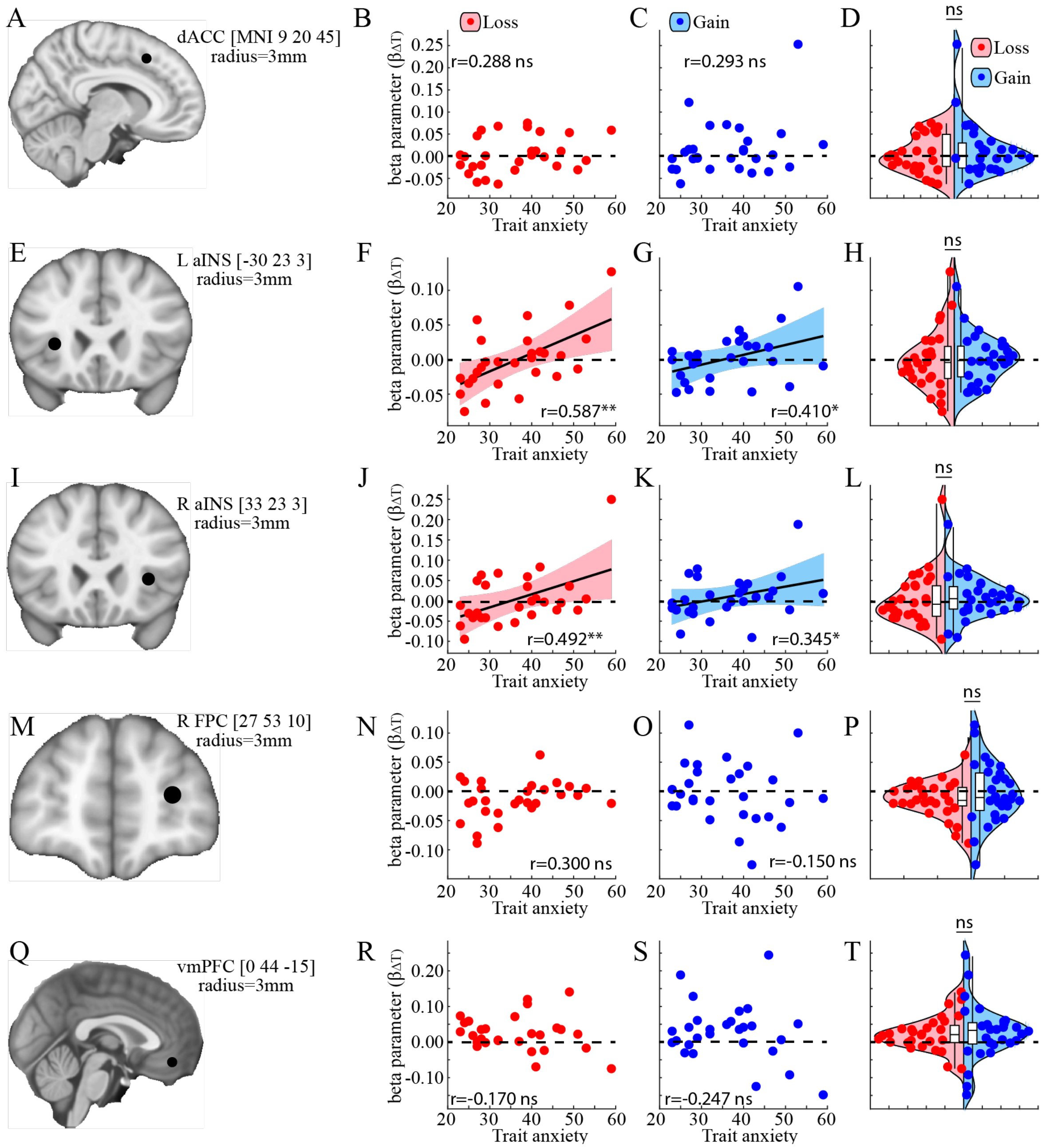

**Supplementary Figure 12.** Beta parameter estimates (i.e. $\beta_{\Delta T}$) of environmental uncertainty (i.e. $\Delta T$) as a function of Gain/Loss condition and Trait anxiety. Beta parameters were extracted from 3mm radius spheres (indicated by black spheres) centered on the coordinates reported in the left-most column. **A-C.** $\beta_{\Delta T}$ did not correlate with trait anxiety in the dACC (**B, C**), nor was there a difference between Gain/Loss conditions (**D**). **E-H.** $\beta_{\Delta T}$ of the left aINS correlated positively with trait anxiety in the Loss (**F**) and in the Gain condition (**G**), but did not differ between Gain/Loss conditions (**H**). Similar results were obtained for the right aINS (**I-J**), while no significant correlations with trait anxiety or differences between Gain/Loss conditions were obtained in the right FPC (**M-P**) or the vmPFC (**Q-T**). **p<0.001; **p<0.01; *p<0.05; ns p>0.05.

**Supplementary Table 7.** Repeated measures ANOVA for $\beta_{\Delta T}$ as a function of Gain/Loss conditions and Trait anxiety in each of the a priori ROIs.

| | Main analysis | | | | | Robustness analysis p-value (uncorrected) | | |
|---|---|---|---|---|---|---|---|---|
| | Sum of Squares | df | Mean Square | F | p-value (uncorrected) r=3 | r=0 | r=5 | r=10 |
| vmPFC | | | | | | | | |
| Intercept | 0.037 | 1 | 0.037 | 6.931 | 0.014 | 0.016 | 0.014 | 0.015 |
| Trait anxiety | 0.011 | 1 | 0.011 | 2.076 | 0.162 | 0.181 | 0.145 | 0.168 |
| Error | 0.137 | 26 | 0.005 | | | | | |
| Condition | <.001 | 1 | <.001 | 0.041 | 0.841 | 0.824 | 0.841 | 0.899 |
| Trait anxiety x Condition | 0.002 | 1 | 0.002 | 0.503 | 0.485 | 0.521 | 0.536 | 0.734 |
| Error | 0.094 | 26 | 0.004 | | | | | |
| dACC | | | | | | | | |
| Intercept | 0.006 | 1 | 0.006 | 2.068 | 0.162 | 0.173 | 0.130 | 0.079 |
| Trait anxiety | 0.012 | 1 | 0.012 | 4.085 | 0.054 | 0.079 | 0.040 | 0.090 |
| Error | 0.076 | 26 | 0.003 | | | | | |
| Condition | 0.002 | 1 | 0.002 | 0.821 | 0.373 | 0.391 | 0.388 | 0.383 |
| Trait anxiety x Condition | <.001 | 1 | <.001 | 0.230 | 0.636 | 0.571 | 0.749 | 0.962 |
| Error | 0.058 | 26 | 0.002 | | | | | |
| Right FPC | | | | | | | | |
| Intercept | <.001 | 1 | <.001 | 3.049 | 0.093 | 0.095 | 0.073 | 0.111 |
| Trait anxiety | <.001 | 1 | <.001 | 0.034 | 0.855 | 0.871 | 0.848 | 0.981 |
| Error | 0.030 | 26 | 0.001 | | | | | |
| Condition | <.001 | 1 | <.001 | 0.690 | 0.414 | 0.445 | 0.389 | 0.328 |
| Trait anxiety x Condition | 0.004 | 1 | 0.004 | 1.596 | 0.218 | 0.214 | 0.207 | 0.223 |
| Error | 0.067 | 26 | 0.003 | | | | | |
| Left aINS | | | | | | | | |
| Intercept | <.001 | 1 | <.001 | 0.006 | 0.873 | 0.873 | 0.947 | 0.709 |
| Trait anxiety | 0.022 | 1 | 0.022 | 15.432 | <.001 * | <.001 * | <.001 * | 0.001 * |
| Error | 0.037 | 26 | 0.001 | | | | | |
| Condition | <.001 | 1 | <.001 | 0.078 | 0.783 | 0.832 | 0.794 | 0.982 |
| Trait anxiety x Condition | 0.002 | 1 | 0.002 | 1.944 | 0.175 | 0.156 | 0.183 | 0.330 |
| Error | 0.025 | 26 | 0.001 | | | | | |
| Right aINS | | | | | | | | |
| Intercept | 0.003 | 1 | 0.003 | 0.876 | 0.360 | 0.360 | 0.438 | 0.664 |
| Trait anxiety | 0.035 | 1 | 0.035 | 11.855 | 0.002 * | 0.001 * | 0.002 * | 0.004 * |
| Error | 0.077 | 26 | 0.003 | | | | | |

| | | | | | | | | |
|---|---|---|---|---|---|---|---|---|
| Condition | <.001 | 1 | <.001 | 0.147 | 0.705 | 0.690 | 0.690 | 0.700 |
| Trait anxiety x Condition | 0.003 | 1 | 0.003 | 0.835 | 0.369 | 0.394 | 0.340 | 0.242 |
| Error | 0.081 | 26 | 0.003 | | | | | |

vmPFC = ventromedial prefrontal cortex. dACC=dorsal anterior cingulate cortex. FPC = right frontopolar cortex. aINS = anterior insula. * indicates that the p-value is significant at the Bonferroni-corrected threshold (α=0.01).

### Supplementary Note 12

To confirm that the relationship between trait anxiety and the behavioral factors that drive exploration, i.e. the tradeoff between expected-value and uncertainty reduction ($\beta_Q - \beta_T$; Figure 4E), is related to the differential representation of expected-value in the dACC (i.e. $\beta_{\Delta Q}$; Figure 6B) and outcome uncertainty in the aINS ($\beta_{\Delta T}$; Figure 6F), a mediation analysis was performed. Specifically, this analysis tests whether the relationship between trait anxiety and $\beta_Q - \beta_T$ is mediated by the differential representation of expected-value in the dACC and outcome uncertainty in the aINS (i.e. $\beta_{\Delta Q,dACC} - \beta_{\Delta T,aINS}$). $\beta_{\Delta T,aINS}$ was calculated across the left and the right aINS. Indeed, the standardized linear regression coefficient between trait anxiety and $\beta_Q - \beta_T$ (bAnxiety1; Supplementary Table 8) was significantly reduced when $\beta_{\Delta Q,dACC} - \beta_{\Delta T,aINS}$ (bAnxiety2; Supplementary Table 8) was added to the regression. Monte-Carlo randomization procedures confirmed significance, both in the Loss (ΔbAnxiety=-0.254, p<0.001; Supplementary Figure 13A, left panel) and in the Gain condition (ΔbAnxiety=-0.292, p<0.001; Supplementary Figure 13A, right panel ).

***Supplementary Table 8. Mediation analysis.***

| | Predictor | $\beta_{Standardized}$±STD | p-value |
|---|---|---|---|
| **Loss condition** | | | |
| Trait anxiety only | $b_{Anxiety1}$ | -0.583 ± 0.159 | 0.001 |
| $\beta_{\Delta Q,dACC} - \beta_{\Delta T,aINS}$ & Trait anxiety | $b_{Brain}$ | 0.434 ± 0.195 | 0.035 |
| | $b_{Anxiety2}$ | -0.329 ± 0.195 | 0.088 |
| | $\Delta b_{Anxiety} = b_{Anxiety1} - b_{Anxiety2}$ | -0.254 | <0.001* |
| | $\Delta b_{Anxiety,\ null}$ | -0.0005 ± 0.035 | |
| **Gain condition** | | | |
| Trait anxiety only | $b_{Anxiety1}$ | -0.593 ± 0.158 | <0.001 |
| $\beta_{\Delta Q,dACC} - \beta_{\Delta T,aINS}$ & Trait anxiety | $b_{Brain}$ | 0.470 ± 0.215 | 0.039 |
| | $b_{Anxiety2}$ | -0.301 ± 0.215 | 0.120 |
| | $\Delta b_{Anxiety} = b_{Anxiety1} - b_{Anxiety2}$ | -0.292 | <0.001* |
| | $\Delta b_{Anxiety,\ null}$ | -0.001 ± 0.038 | |

Trait anxiety only: only trait anxiety was used as a linear predictor of $\beta_Q - \beta_T$. $\beta_{\Delta Q,dACC} - \beta_{\Delta T,aINS}$ & Trait anxiety: both trait anxiety and $\beta_{\Delta Q,dACC} - \beta_{\Delta T,aINS}$ were used as linear predictors of $\beta_Q - \beta_T$. bAnxiety denotes the standardized predictive value of Trait anxiety, while bBrain denotes the standardized predictive value of $\beta_{\Delta Q,dACC} - \beta_{\Delta T,aINS}$. ΔbAnxiety denotes the difference in predictive value of Trait anxiety without and with $\beta_{\Delta Q,dACC} - \beta_{\Delta T,aINS}$, and its statistical significance was determined using a Monte-Carlo randomization procedure. ΔbAnxiety, null denotes the null-distribution obtained from the Monte-Carlo randomization procedure. STD is the standard deviation of the mean. * Statistically significant effect of mediation, i.e. p-value<0.05.

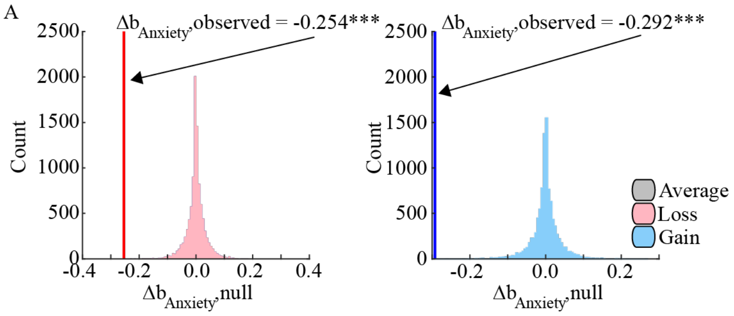

**Supplementary Figure 13.** Change in the linear relationship between Trait anxiety and the tradeoff between expected-value and uncertainty-reduction ($\beta_Q - \beta_T$) when controlling for the difference in the neuronal representation of expected-value in the dACC and outcome uncertainty in the aINS ($\beta_{\Delta Q,dACC} - \beta_{\Delta T,aINS}$). The observed difference in ΔbAnxiety (red line) differed significantly from the null-distribution of differences ΔbAnxiety, null (blue bars) in the Loss condition **(A)** and in the Gain condition **(B)**. ***p<0.001.
